# Proteomic mapping of the dengue virus NS1 microenvironment in infected cells identifies novel host dependency factors including TM9SF3

**DOI:** 10.64898/2026.07.22.740010

**Authors:** Siena Centofanti, Alex Colella, Nusha Chegeni, Kamelya Aliakbari, Gustavo Bracho, Eva Hesping, Michael Roach, Jillian M. Carr, Tim Chataway, Nicholas S. Eyre

**Affiliations:** College of Medicine and Public Health, Flinders University, Bedford Park, Adelaide, South Australia, Australia; Flinders Proteomics Facility, College of Medicine and Public Health, Flinders University, Bedford Park, Adelaide, South Australia, Australia; Cell Screen SA, College of Medicine and Public Health, Flinders University, Bedford Park, Adelaide, South Australia, Australia

## Abstract

Dengue virus (DENV) is endemic in over 100 countries and causes approximately 100 million symptomatic infections annually, with symptoms ranging from mild febrile illness to life-threatening severe vascular leakage and haemorrhagic fever. There are currently no approved antiviral therapies available to treat DENV infections. The DENV non-structural protein 1 (NS1) is essential for viral RNA replication and infectious virus particle production, while secreted NS1 contributes to immune evasion and pathogenicity. Towards the identification of novel NS1-host protein interactions that are critical to these functions, an APEX2 proximity labelling-coupled quantitative proteomics approach was employed to map the proteomic composition of the NS1 microenvironment in live infected cells. Our analysis identified a panel of 51 NS1-proximal host proteins, including established DENV host dependency factors (HDFs) involved in NS1 folding and *N*-glycosylation, as well as previously unrecognised host factors. Loss-of-function approaches were used to determine the importance of these NS1-proximal host proteins to DENV infection, identifying several novel HDFs, including transmembrane 9 superfamily member 3 (TM9SF3). Importantly, the knockout of TM9SF3 was shown to impair DENV infectious virion production and intracellular NS1 abundance and secretion, consistent with the recently described roles of TM9SF3 in Golgi integrity and glycosylation fidelity. Together, this study demonstrates the successful application of APEX2 proximity labelling-coupled quantitative proteomics to the identification of functionally relevant NS1-associated host proteins that may inform the development of future antiviral therapies.

**IMPORTANCE:** The DENV NS1 protein is a non-enzymatic, multifunctional glycoprotein that plays multiple distinct roles in viral replication organelle formation, viral RNA replication and infectious virus particle production. It is also secreted from infected cells as an oligomeric lipoparticle that participates in immune evasion and vascular damage. Many of its enigmatic roles are thought to be mediated via its interactions with other viral proteins and host proteins. Here, we have employed an infectious NS1-tagged DENV reporter virus and proximity biotinylation-coupled mass spectrometry to characterise the protein microenvironment of NS1 during viral infection. We have then employed functional genomics approaches to identify NS1-proximal host factors that contribute to the viral replication cycle. Amongst the novel host factors that were identified was TM9SF3, which has recently emerged as a Golgi-resident Golgiphagy receptor that is important for maintenance of Golgi integrity and glycosylation fidelity and may represent a future DENV antiviral drug target.

## INTRODUCTION

Dengue virus (DENV) is the most prevalent mosquito-borne human pathogen, causing an estimated 400 million infections annually across tropical and subtropical regions worldwide, of which approximately 100 million are symptomatic (1). More than half of the world’s population live in areas that are suitable for DENV transmission (2). Infections lead to a broad spectrum of clinical outcomes, ranging from self-limiting febrile illness to life-threatening disease characterised by plasma leakage, haemorrhaging and shock. There are currently no approved antiviral therapies for dengue infections, highlighting the need to better understand virus-host interactions that could serve as novel antiviral drug targets.

DENV belongs to the *Orthoflavivirus* genus within the *Flaviviridae* family. Following entry into susceptible host cells, the 11 kb positive-sense single-stranded viral RNA genome is released into the cytoplasm, where it is translated into a single polyprotein by host cell machinery. The polyprotein is cleaved by viral and host proteases into three structural proteins (capsid [C], envelope [E] and premembrane [prM]) and seven non-structural (NS) proteins (NS1, NS2A, NS2B, NS3, NS4A, NS4B and NS5). Viral RNA replication occurs within specialised vesicles derived from virus-induced rearrangement of endoplasmic reticulum membranes, in which non-structural proteins assemble to form the viral replication complex (RC). Newly synthesised viral genomes associate with capsid proteins to form nucleocapsids, which bud into the ER lumen to acquire a membrane embedded with E and prM proteins. These immature virions undergo maturation through furin-mediated cleavage of prM during transit through the Golgi before being secreted from the cell (3).

DENV NS1 is a 48-kDa glycoprotein critical to multiple stages of the viral lifecycle. Following cleavage from the viral polypeptide and translocation into the ER lumen, NS1 monomers are glycosylated at two conserved *N*-linked glycosylation sites, N130 and N207, with high-mannose glycans (4–6). Glycosylation at these sites is critical to the stability of the oligomers that are subsequently formed by NS1 and is important for the efficient secretion of NS1 (7, 8). NS1 monomers rapidly dimerise, which imparts partial hydrophobicity, enabling stable association with ER-derived virus-induced replication organelles (4, 9, 10). During transit through the Golgi, the N130-linked glycan is trimmed and further modified into a complex multi-branched form (6). NS1 dimers can form soluble tetramers or hexamers, the latter of which is stabilised by a lipid-rich core, and are secreted from infected cells (6, 11–14). It is unclear whether the formation of NS1 hexameric lipoparticles occurs within the ER following dimerization, or during transit through the Golgi (15). The mechanistic understanding of intracellular NS1 functions has largely been informed by mutagenesis studies. These studies have shown that NS1 performs distinct and indispensable roles in the biogenesis of viral replication organelles, facilitating viral RNA synthesis via its interaction with the NS4A-2K-4B cleavage intermediate, and the production of infectious virus particles, potentially mediated through interactions with the structural proteins E and prM (16, 17). Secreted NS1 is a key pathogenic factor that induces vascular hyperpermeability by triggering pro-inflammatory cytokine release and disrupting the endothelial glycocalyx and intracellular junctions (18–22). Additionally, secreted NS1 contributes to the evasion of host immune responses by antagonising and supressing complement activation (23–25). Circulating levels of secreted NS1 in patients correlate with viremia and greater disease severity (26). Given the essential roles of NS1 in viral replication and pathogenesis, defining the interactions of NS1 that support these functions may uncover novel therapeutic drug targets.

In this study, a novel infectious APEX2-tagged DENV reporter virus together with APEX2 proximity labelling-coupled quantitative proteomics was employed to map the proteomic composition of the NS1 protein microenvironment during a productive infection. APEX2 is a monomeric 28-kDa engineered ascorbate peroxidase tag that is fused either to a signal peptide, enabling the mapping of proteomes within membrane-bound cellular compartments, or to a protein of interest, allowing the identification of the proteomic composition of its surrounding microenvironment (27). APEX2 enables the spatially restricted tagging of proteins in live cells by catalysing the hydrogen peroxide (H2O2)- dependent oxidation of biotin-phenol (BP) into short-lived, membrane-impermeant BP radicals during a brief (≤ 1 minute) labelling period. The BP radicals covalently tag the electron-rich amino acids of proteins that are proximal to APEX2, resulting in their biotinylation. While it is estimated that APEX2 labelling radius is 10-20 nm, labelling occurs across a gradient, with proteins closest to APEX2 being more heavily biotinylated than distal proteins. The biotinylated proteins can then be enriched by streptavidin pulldown, digested with trypsin and identified by mass spectrometry.

The functional relevance of the identified NS1-proximal host proteins was interrogated using a targeted siRNA screen and CRISPR-Cas9-mediated gene silencing. This resulted in the identification of both well-characterised and previously unrecognised DENV HDFs required for efficient viral infection. Further functional characterisation revealed an important role for TM9SF3, a transmembrane protein critical for Golgi integrity and glycosylation homeostasis, in DENV infectious virion production and NS1 abundance and secretion. Our study provides the first proximity labelling-coupled quantitative proteomics-based map of host proteins associated with DENV NS1 in live infected cells and demonstrates the utility of this approach for identifying novel host dependency factors and host pathways that support the DENV lifecycle and NS1 biology.

## RESULTS

### Generation and Characterisation of APEX2-Tagged Reporter Viruses

To label proteins that are proximal to NS1 during a productive DENV infection, we utilised an infectious DENV2-NS1-APEX2 reporter virus which encodes an APEX2 tag within the second connector domain of NS1 in a full-length DENV2 cDNA clone (Fig. 1A) (10). In a previous study, this reporter virus was characterised and employed for APEX2-mediated 3,3’-diaminobenzidine (DAB) labelling and electron microscopy analysis, revealing expected localization of NS1 to vesicle packets (VPs) and, to a lesser extent, Golgi stacks and highly ordered arrays (10). In the present study we sought to employ this reporter virus towards proximity biotinylation-coupled mass spectrometry to identify NS1 proximal proteins in the context of a productive DENV infection (Fig. 1B). In the context of proximity biotinylation, the greatest degree of protein biotinylation occurs within the immediate vicinity of APEX2, due to the short lifetime and limited diffusion of BP radicals (27). However, labelling can extend beyond this range if the protein of interest (POI)-APEX2 fusion is not contained within organelle membranes and, while unlikely, ‘off-target’ labelling can result from unanticipated interactions of the APEX2 protein itself. To control for off-target labelling in the context of a DENV2-infected cell and with APEX2 expression levels that are comparable to those of viral proteins, a control reporter virus (DENV2-2A-APEX2-2A) was generated by inserting APEX2 flanked by 2A self-cleaving peptides immediately downstream of the capsid leader sequence (Fig. 1A). Inclusion of the 2A self-cleaving peptides results in the co-translational cleavage of APEX2 from the viral polypeptide, allowing the tag to diffuse freely within the cell and catalyse non-targeted protein biotinylation, whilst being co-expressed with viral proteins.

**Figure 1.**
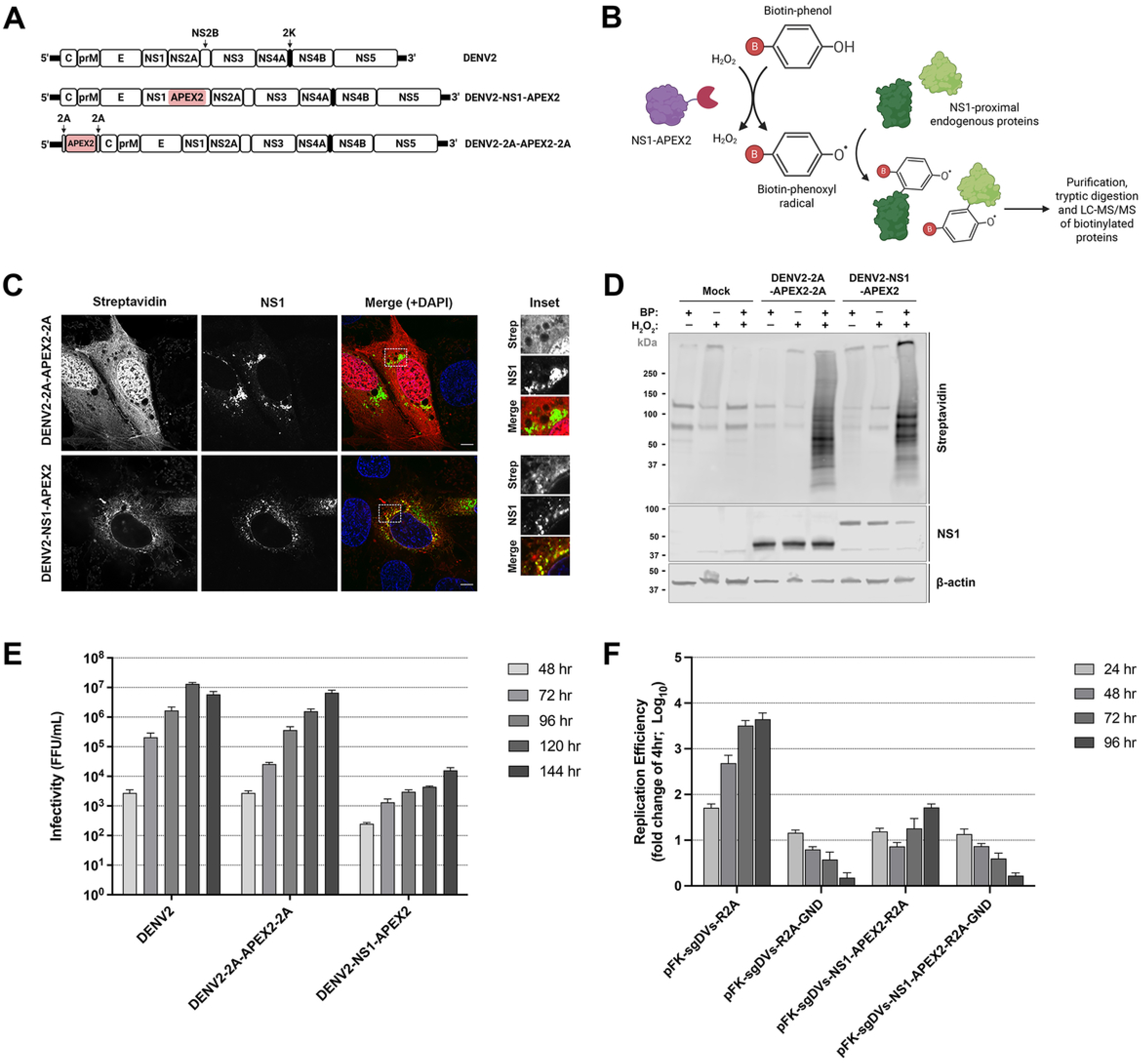
Design and characterization of DENV2 reporter viruses encoding an APEX2 tag. (A) Schematic of the full-length DENV2 constructs used in this study. (B) Schematic of the APEX2-mediated biotinylation of NS1-proximal endogenous proteins. (C) Localisation of NS1 with respect to biotinylated proteins within Huh-7.5 cells transfected with DENV2-2A-APEX2-2A or DENV2-NS1-APEX2 IVT RNA. Cells were transfected with the indicated DENV2 RNA transcripts and treated with BP/H2O2 at 5 days post-transfection prior to fixation and immunofluorescent labelling using anti-NS1, Alexa Fluor 488-conjugated anti-mouse IgG (green) and streptavidin-conjugated Cy3 (red). Samples were stained with DAPI (blue) and analyzed by confocal fluorescence microscopy. Yellow in the merged channels is indicative of colocalization. Insets are zoomed-in images of the boxed areas in the merged channels. Scale bars represent 5 µm. (D) Streptavidin blot analysis of biotinylated proteins within lysates collected from Huh-7.5 cells transfected with DENV2-2A-APEX2-2A or DENV2-NS1-APEX2 RNA. Cells were transfected with the indicated DENV2 RNA transcripts and treated with BP/H2O2 at 5 days post-transfection prior to lysis. Total proteins from lysed cells were separated by SDS-PAGE and analysed by Western blotting using fluorescently labelled streptavidin, anti-NS1 and anti-β-actin. (E) Infectious virus production in Huh-7.5 cells transfected with DENV2, DENV2-2A-APEX2-2A or DENV2-NS1-APEX2 IVT RNA. Cells were transfected with the indicated IVT DENV2 RNA transcripts and returned to culture prior to the collection of supernatants at the indicated time points (48 to 144 hours). Infectivity of virus within supernatant samples was determined by focus-forming assay. Data are presented as means ± the standard deviations (SD; n = 3 independent replicates). (F) Replication efficiency in Huh-7.5 cells transfected with sgDVS-R2A or sgDVS-NS1-APEX2-R2A. Cells were transfected with the indicated IVT DENV2 subgenomic replicon RNA transcripts and lysates were harvested at 4, 24, 48, 72 and 96 hours post-transfection for measurement of *Renilla* luciferase activity, an indicator of viral RNA replication efficiency. For each subgenomic replicon, raw luciferase values were normalised to those of the 4-hour timepoint average value (representative of input RNA) to account for differences in transfection efficiencies. Replication deficient mutants harbouring a ‘GND’ mutation within NS5 were included as negative controls. Data are presented as means ± S.D. (n = 3).

To characterise DENV2-2A-APEX2-2A- and DENV2-NS1-APEX2-mediated protein biotinylation, Huh-7.5 hepatoma cells were transfected with *in vitro* transcribed (IVT) DENV2-NS1-APEX2 and DENV2-2A-APEX2-2A RNA transcripts and treated with BP and H2O2 at 5 days post-transfection. The cells were then lysed for Western blotting analysis or fixed for indirect immunofluorescent staining and confocal imaging. Strong co-localisation between biotinylated proteins and NS1 within cells transfected with IVT DENV2-NS1-APEX2 RNA indicates that this reporter virus enables the biotinylation of proteins that are within the NS1 microenvironment during a productive infection (Fig. 1C). Co-localization of biotinylated and NS1 proteins was primarily observed in the perinuclear region of the cells. As expected, biotinylated protein-associated signals were seen to also extend beyond NS1 foci, with previous studies demonstrating that biotinylated proteins migrate from APEX2 prior to cell lysis or fixation (28). In contrast, biotinylated proteins were widely distributed throughout the nuclear, peri-nuclear and cytosolic regions of cells transfected with IVT DENV2-2A-APEX2- 2A RNA, suggesting that APEX2 was efficiently cleaved from the DENV2 polyprotein and migrated freely throughout the cell, resulting in non-targeted protein biotinylation (Fig. 1C). Biotinylated protein-associated signals were notably stronger in these cells compared to those of cells that were transfected with IVT RNA for DENV2-NS1-APEX2. Streptavidin blotting of biotinylated proteins revealed an abundance of biotinylated proteins across a large range of molecular weights for DENV2-NS1-APEX2 and DENV2-2A-APEX2-2A IVT RNA-transfected cells treated with BP and H2O2 compared to negative controls for which APEX2, BP or H2O2 was omitted, indicating that the virally encoded APEX2 proteins are catalytically active (Fig. 1D). Distinct banding patterns were detected for DENV2-NS1- APEX2 and DENV2-2A-APEX2-2A IVT RNA-transfected cells, reflecting differential subcellular localisation and targeting of APEX2-mediated biotinylation for these viruses. As expected, the NS1-APEX2 fusion protein (76 kDa) and NS1 protein (48 kDa) were readily detected in lysates from DENV2-NS1-APEX2- and DENV2-2A-APEX2-2A-infected cells, respectively, with NS1-APEX2 bands appearing weaker than those of untagged NS1 (Fig. 1D). Weak bands at ∼130 kDa and ∼75 kDa were detected by streptavidin blotting for all samples including the negative controls and correspond to similar bands that have been reported for endogenously biotinylated proteins expressed by mammalian cells (28, 29).

Importantly, we have previously shown by Western blotting that NS1 expression, secretion and *N*-glycosylation in Huh-7.5 cells transfected with DENV2-NS1-APEX2 IVT RNA appears comparable to that of cells transfected with DENV2 IVT RNA (10). Furthermore, in cells transfected with DENV2-NS1-APEX2 IVT RNA, the subcellular localisation of NS1 appeared unaltered, with only very minor changes observed in its colocalization with dsRNA, a marker of viral replication complexes (10). Following these observations, we sought to further characterise the impact of APEX2 insertion on DENV2 replication and infectious virus particle production. Huh-7.5 cells were transfected with DENV2, DENV2-2A-APEX2-2A or DENV2-NS1-APEX2 IVT RNA, and culture supernatants were harvested at 48-144 hours post-transfection for measurement of viral infectivity. Compared to wildtype DENV2, DENV2-NS1-APEX2 showed appreciable attenuation at all time points, with a ∼2.6-log reduction in infectious virus production at 144 hours post-transfection, whereas DENV2-2A-APEX2-2A maintained near-wildtype levels of infectious virus production at all timepoints (Fig. 1E). To evaluate the impact of the APEX2 insertion on viral RNA replication, independent of infectious virus production, the APEX2-tagged NS1 gene was incorporated into a *Renilla* luciferase-encoding DENV2 subgenomic replicon construct (pFK-sgDVs-R2A). Huh-7.5 cells were transfected with IVT RNA for sgDVs-NS1- APEX2-R2A, sgDVs-R2A or replication-defective ‘GND’ mutants of these replicons. Lysates were harvested at various timepoints post-transfection for measurement of luciferase activity, which reflects levels of viral translation and RNA replication. Whilst the replication of sgDVs-NS1-APEX2-R2A steadily increased across all time timepoints, it was appreciably attenuated compared to the wildtype subgenomic replicon, displaying a ∼1.9-log reduction in replication efficiency at 96 hours post-transfection (Fig. 1F). Together, these results indicate that the insertion of APEX2 within NS1 significantly attenuates DENV2 replication and infectious virus production without appreciably altering NS1 secretion, glycosylation or subcellular localization.

### Proteomic Mapping of the NS1 Microenvironment during a Productive DENV Infection

To map the proteomic composition of the NS1 microenvironment, we utilized the DENV2- NS1-APEX2 and DENV2-2A-APEX2-2A reporter viruses in combination with an APEX2 proximity labelling-coupled three-state SILAC quantitative proteomics approach adopted from Hung *et al.* (Fig. 2A) (28). Huh-7.5 cells were cultured in SILAC media supplemented with heavy, medium or light isotope-labelled amino acids for 7 cell doublings and then transfected with DENV2-NS1-APEX2 or DENV2-2A-APEX2-2A IVT RNA, or mock-transfected. To control for potential impacts of SILAC labelling conditions on mass spectrometry readouts, the RNA transfection conditions were alternated between the SILAC labelling conditions across three independent experimental replicates (Fig. 2B). At 5 days post-transfection, the cells were incubated with BP and then briefly treated with H2O2 to initiate APEX2-mediated protein biotinylation. The reactions were quenched, and lysates were harvested from the heavy, medium and light SILAC-labelled cell populations. Streptavidin blotting confirmed robust biotinylation in DENV2-NS1-APEX2 and DENV2-2A- APEX2-2A IVT RNA-transfected cells (Fig. S1). As expected, distinct banding patterns were observed for DENV2-NS1-APEX2 and DENV2-2A-APEX2-2A samples, and these patterns were reproducible across the three independent experimental replicates. Importantly, analysis of the degree of SILAC labelling revealed near-complete saturation of heavy, medium and light protein labelling in corresponding experimental groups (Tables S1 and S2). Lysates from the heavy, medium and light SILAC-labelled cell populations were pooled in a 1:1:1 ratio based on protein mass and biotinylated proteins were enriched by streptavidin pulldown, separated by SDS-PAGE and digested with trypsin (Fig. S2). The identity and relative abundance of proteins each of heavy, medium and light SILAC-labelled cell population were determined by liquid chromatography tandem mass spectrometry (LC-MS/MS) (Table S3). Pairwise comparisons between biological replicates of each experimental condition showed a high correlation (R² = 0.85–0.89) in SILAC-based measurements of protein abundance (Fig. S3). The proteins that were significantly enriched in DENV2-NS1-APEX2 samples relative to both DENV2-2A-APEX2 and mock samples (log2 fold change > 1, adjusted *p*-value < 0.05) were considered NS1-proximal (Fig. 2C and S4).

**Figure 2.**
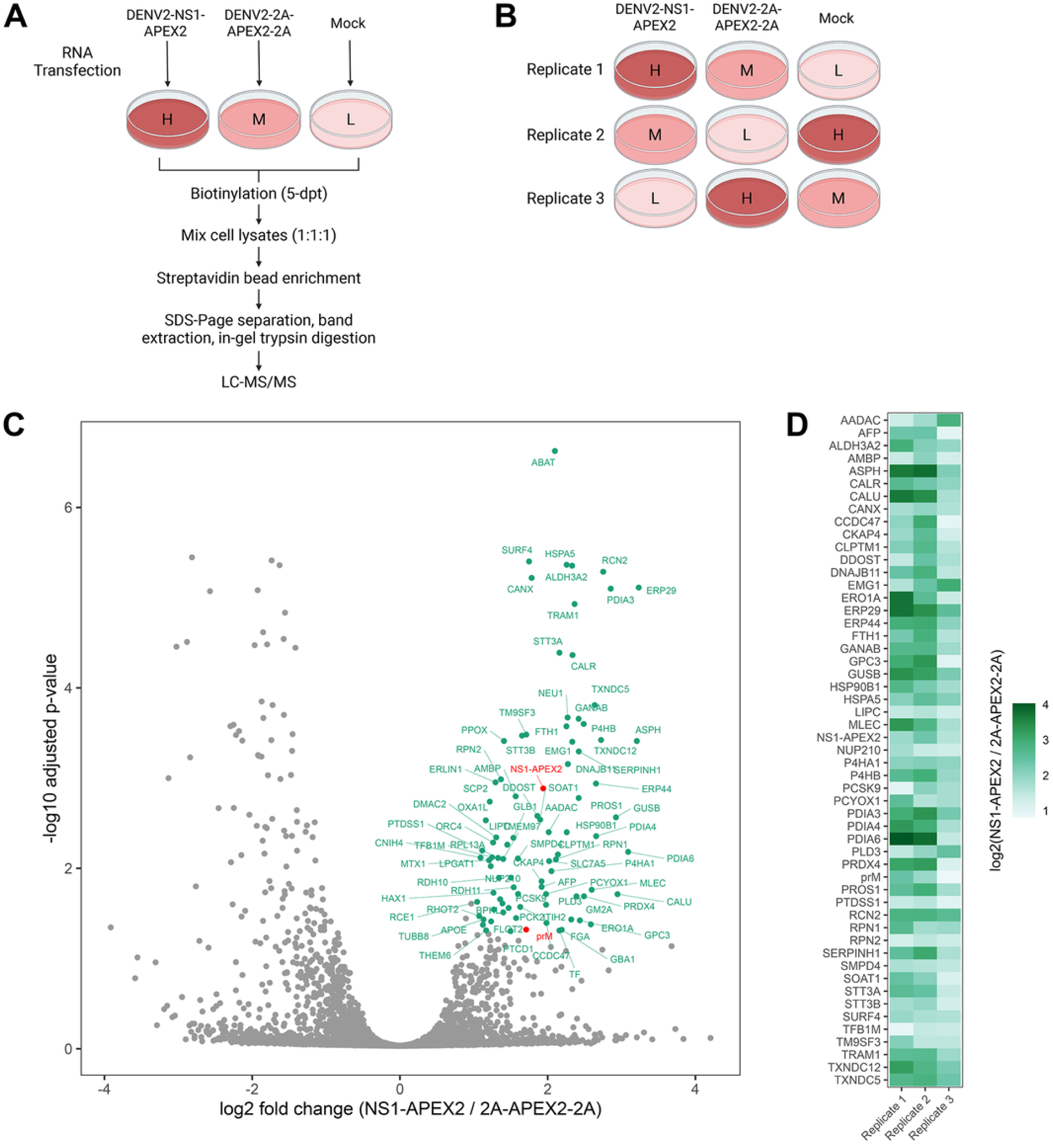
Proteomic analysis of biotinylated NS1-proximal proteins. (A) Schematic of the APEX2 proximity labelling-coupled quantitative proteomics strategy used to identify NS1-proximal proteins. (B) Schematic of the SILAC labelling strategy. Heavy (H), medium (M) and light (L) labelling conditions were rotated between DENV2-NS1-APEX2, DENV2-2A-APEX2-2A and mock-transfected controls across three independent experiments. (C) Volcano plot of the relative enrichment of proteins in DENV2-NS1-APEX2 samples compared with DENV2-2A-APEX2-2A samples. Proteins with log2 fold change > 1 and adjusted p-value < 0.05 are considered significantly enriched in DENV2-NS1-APEX2 samples and are highlighted in green (host proteins) or red (DENV2 proteins). The depicted plot represents a merged dataset from three independent experiments. (D) Heat map of log₂-transformed abundance ratios (NS1-APEX2 / 2A-APEX2-2A) across three independent experiments for proteins significantly enriched in DENV2-NS1-APEX2 samples relative to the DENV2-2A-APEX2-2A and mock samples.

A total of 53 proteins were identified as NS1-proximal, including 51 host proteins and 2 viral proteins, prM and NS1-APEX2 (Fig. 2D). Gene Ontology analysis revealed a strong enrichment of ER-resident host proteins involved in protein folding and glycoprotein maturation, including the chaperones CALR, CANX, HSPA5 and HSP90B1, glucosidase II subunit GANAB and oligosaccharyltransferase (OST) complex subunits DDOST, RPN1, RPN2, STT3A and STT3B (Fig. 3A and 3B). Protein disulfide isomerases (P4HB, PDIA3, PDIA4, PDIA6, TXNDC5, TXNDC12), which catalyse formation, reduction and isomerisation of disulfide bonds to promote redox protein folding and protein quality control, were significantly enriched. Several oxidoreductases (ERO1A, ERP29, ERP44 and PRDX4), which catalyse redox reactions to maintain ER oxidative homeostasis and support protein disulfide isomerase activity, were also identified. ER-resident co-chaperones (DNAJB11, ERP29 and ERP44), calcium-binding proteins (CALR, CALU and RCN2), and regulators of lipid metabolism and cholesterol homeostasis (LIPC, PCSK9, PCYOX1, PLD3, PTDSS1, SMPD4, SOAT1) were also enriched. Of the 51 NS1-proximal host proteins, 14 have previously been identified as DENV host dependency factors (Table S4). Collectively, our results reveal that the NS1 microenvironment is highly enriched for ER-resident chaperones, enzymes that catalyse post-translational protein modifications, and lipid regulators, consistent with the dependence of NS1 on the secretory pathway for correct folding, maturation and trafficking as a secreted glycoprotein. However, a substantial fraction of the identified host factors represent novel NS1-associated proteins and highlight potential new host dependencies for DENV2.

**Figure 3.**
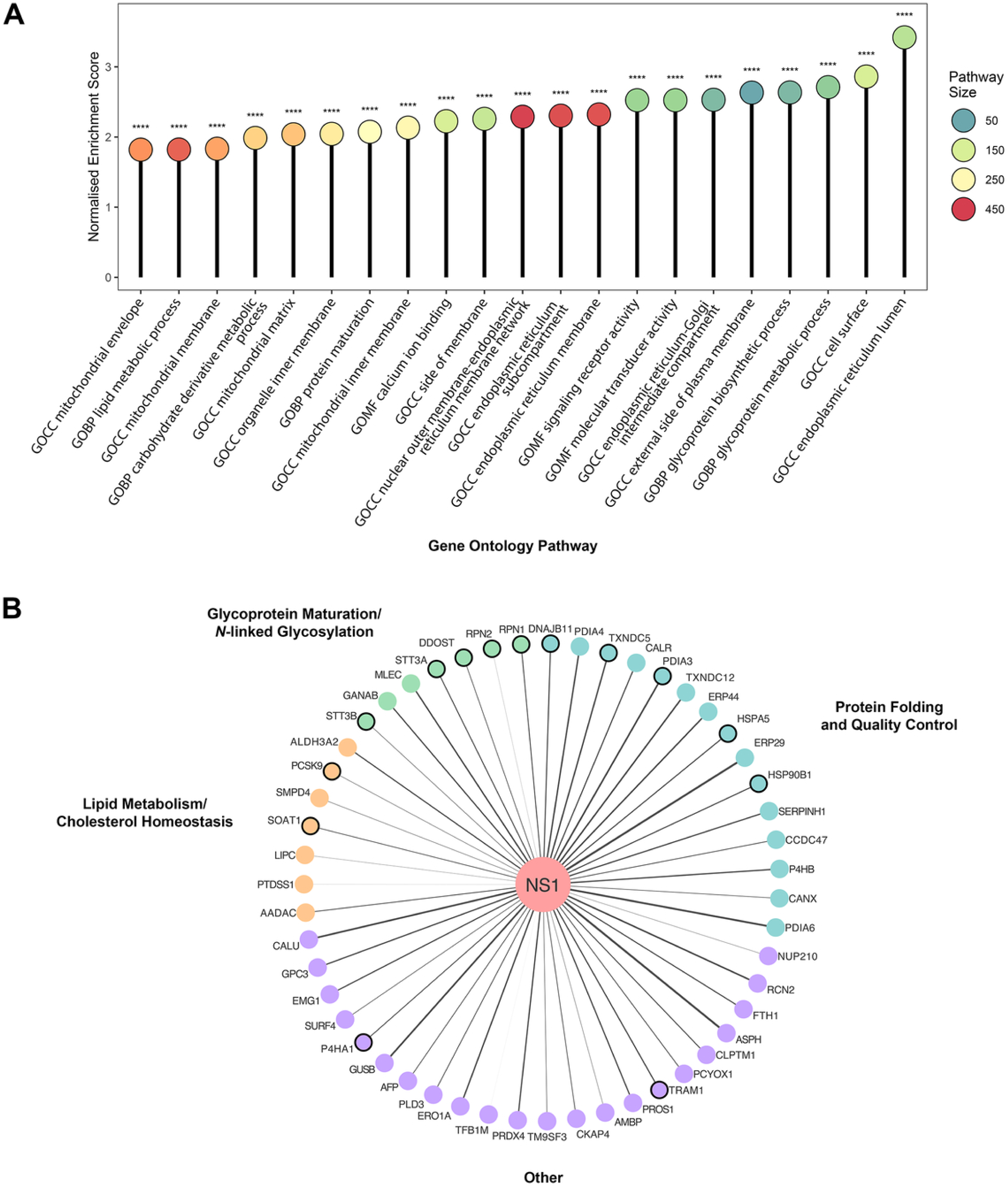
Gene ontology (GO) analysis of biotinylated NS1-proximal proteins. (A) Gene ontology (GO) enrichment analysis of biological processes (BP), molecular functions (MF) and cellular compartments (CC) for proteins significantly enriched in DENV2-NS1-APEX2 samples relative to the DENV2-2A-APEX2-2A and mock samples. Pathways are ranked by normalised enrichment score. Circle colour indicates pathway size and statistical significance is indicated by adjusted p-values (****p<0.0001). (B) Network showing the 51 host proteins identified as being proximal to NS1. Edge weights reflect the enrichment score in DENV2-NS1-APEX2 samples relative to the DENV2-2A-APEX2-2A samples. Host proteins previously identified as DENV host dependency factors are indicated by a node border.

### Identification of NS1-Proximal Host Factors that are Essential for DENV Infection

To identify NS1-proximal host factors that are critical to the DENV replication cycle, we utilised an infectious DENV2-NS1-mScarlet reporter virus to determine the effect of silencing select host factors on viral infection. The DENV2-NS1-mScarlet reporter virus encodes the mScarlet fluorescent protein within the NS1 gene of pFK-DVs, enabling the direct visualization of viral infection levels and NS1 protein localization and traffic in live infected cells (Fig. 4A) (10). Additionally, this reporter virus harbours several adaptive mutations that restore viral replicative fitness to near-wildtype levels (Centofanti, Johnson and Eyre; unpublished). A schematic overview of the siRNA screen strategy is provided in Figure 4B. Briefly, Huh-7.5 cells were reverse-transfected with a custom siRNA library of 40 siRNA pools, each consisting of four individual siRNA duplexes that recognise different sequences within a target gene. The 40 host genes that were targeted by these siRNA pools included a selection of 32 host factors that were definitively identified in the proteomics analysis as NS1-proximal proteins and 8 additional host factors of interest that approached but did not meet hit criteria. At 20 hours post-transfection, the cells were infected with DENV2-NS1-mScarlet and cultured for 48 hours before fixation, staining of nuclei and automated fluorescence microscopy. For each siRNA treatment, viral infection was quantified as the percentage of NS1-mScarlet-positive cells, and the percentage of inhibition calculated relative to that of cells transfected with the non-targeting control (NTC) siRNA (Fig. 4C, 4D and S5).

**Figure 4.**
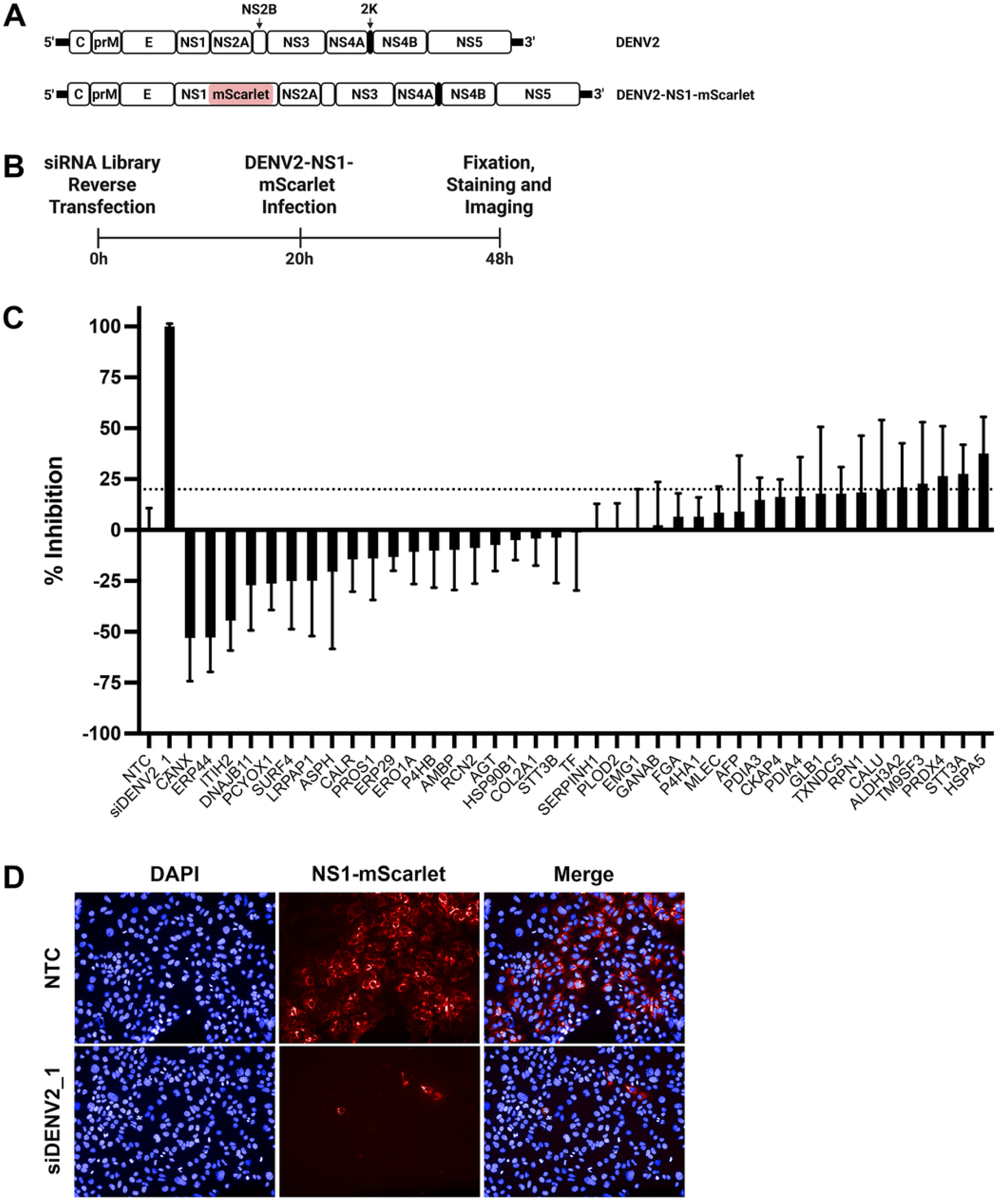
An siRNA screen identifies NS1-proximal host factors that are important for efficient DENV infection. (A) Schematic of the full-length DENV2-NS1-mScarlet construct used in the siRNA screen. (B) Schematic of the siRNA screen. (C) Percent inhibition of DENV2-NS1-mScarlet infection by siRNAs targeting individual NS1-proximal host factors, normalised to those of the non-targeting control (NTC). The host proteins targeted by siRNA treatments that inhibited infection by more than 20% (CALU, ALDH3A2, TM9SF3, PRDX4, STT3A and HSPA5) were considered as pro-viral host proteins required for efficient DENV2 infection. Data are medians ± the median absolute deviation (MAD) from 12 measurements from three independent experiments. (D) Representative images of DENV2-NS1-mScarlet infection in cells transfected with the NTC or siDENV2_1 positive control siRNA.

The siRNA-mediated silencing of six NS1-proximal host factors inhibited DENV infection by ≥20%, including CALU (20.0% inhibition), ALDH3A2 (21.1% inhibition), TM9SF3 (22.8% inhibition), PRDX4 (26.5% inhibition), STT3A (27.5% inhibition) and HSPA5 (37.6% inhibition). Both STT3A and HSPA5 are known orthoflaviviral HDFs and have been shown to be critical to the *N*-linked glycosylation and the folding and secretion of NS1, respectively (30–32). The remaining 4 host proteins identified as hits in our screen represent novel host dependency factors. These include ALDH3A2, an aldehyde dehydrogenase that oxidises medium- and long-chain fatty aldehydes into fatty acids; CALU, a calcium-binding ER chaperone that regulates vitamin K–dependent γ-carboxylation of proteins and contributes to calcium homeostasis; PRDX4, an ER peroxiredoxin that reduces hydrogen peroxide and organic peroxides, contributing to redox homeostasis, protection against oxidative stress and disulphide bond formation during protein folding; and TM9SF3, a transmembrane protein that functions as a Golgiphagy receptor critical to Golgi morphology and glycosylation fidelity (33–38). Several siRNA treatments increased DENV infection levels relative to that of the NTC siRNA, indicating that depletion of these host factors enhanced viral infection (Fig. 4C). As the data are presented as percent inhibition relative to the NTC control, these increases are represented as negative inhibition values. The most pronounced effects were observed for the siRNA-mediated knockdown of CANX (-53.1% inhibition), ERP44 (-52.8% inhibition), ITIH2 (-44.5% inhibition), DNAJB11 (-27.2% inhibition), PCYOX1 (-26.3% inhibition), SURF4 (-25% inhibition), LRPAP1 (-24.9% inhibition) and ASPH (-20.5% inhibition). These proteins are primarily involved in ER protein folding, post-translational modification, quality control, trafficking, or extracellular matrix regulation. Taken together, these results support the validity of the APEX2 proximity labelling-coupled quantitative proteomics approach to capture known host dependency factors that are critical for DENV infection, as well as novel host factors that warrant further investigation.

### TM9SF3 Knockout Impairs DENV Infection, Infectious Virion Production and Intracellular NS1 Protein Abundance and Secretion

To validate the functional relevance of the 6 NS1-proximal host factors identified by siRNA screening as important for DENV infection, CRISPR/Cas9-knockout Huh-7.5 cell lines were generated for each gene and assessed for their ability to support DENV infection. Parental and knockout Huh-7.5 cells were infected with DENV2-NS1-mScarlet and infection levels were monitored by live-cell imaging over a 48-hour period. Across all time points, a 60% or greater reduction in infection levels was observed in one STT3A knockout cell line and in both TM9SF3 knockout cell lines (Fig. 5A). At 48 hours post-infection, relative to parent cells (set at 100%), 10% of STT3A (gRNA 2) knockout cells, 28% TM9SF3 (gRNA 1) knockout cells and 40% of TM9SF3 (gRNA 2) knockout cells were infected (Fig. 5A). As the role of STT3A in the orthoflaviviral lifecycle has been explored in several previous studies, the present study focused on TM9SF3, which has not previously been identified as a host-dependency factor (30, 39–41). To determine the impact of CRISPR-Cas9-medated TM9SF3 knockout on DENV RNA replication, parent and TM9SF3-knockout (gRNA 1) Huh-7.5 cells were transfected with IVT sgDVs-R2A or sgDVs-R2A-GND RNA and lysates were harvested at various timepoints post-transfection for measurement of luciferase activity. The replication efficiency of sgDVs-R2A was comparable in parent and TM9SF3-knockout cells at all time points (Fig. 5B). Following this observation, parent and TM9SF3-knockout (gRNA 1) Huh-7.5 cells were infected with DENV (MOI ∼0.1) and cell supernatants were harvested at 48 and 72 hours post-infection for measurement of viral infectivity. At 48 hours post-infection, a ∼0.75-log reduction in infectious virus titre was observed in TM9SF3 knockout cell supernatants compared to those of parent cells, although this difference was not statistically significant (unpaired *t*-test; *p* > 0.05) (Fig. 5C). However, at 72 hours post-infection, the infectious virus titre was significantly reduced by ∼0.71-log in TM9SF3 knockout cell supernatants compared to those of parent cells (Fig. 5C). Taken together, these results indicate that TM9SF3 is dispensable for viral RNA replication but may be required for efficient production of infectious virions. We next assessed the impact of TM9SF3 knockout on intracellular NS1 protein abundance and secretion. Parent and TM9SF3-knockout Huh-7.5 cells were infected with DENV (MOI ∼0.1), and cell culture supernatants and lysates were harvested at 48 hours post-infection for the detection of secreted and intracellular NS1, respectively, by quantitative Western blot analysis. In TM9SF3-knockout cells, intracellular NS1 abundance was reduced by ∼77% (gRNA 1) and ∼52% (gRNA 2) relative to parent cells (Fig. 5D and 5E). NS1 secretion efficiency (secreted NS1 / intracellular NS1), expressed relative to that of the parent cells, was reduced by ∼48% (gRNA 1) and ∼46% (gRNA 2) in TM9SF3-knockout cells (Fig. 5D and 5F). Given the effects of TM9SF3 knockout on intracellular NS1 abundance and secretion, we next examined NS1 and TM9SF3 localization. Huh-7.5 cells were infected with DENV (MOI ∼0.1) and transfected with a TM9SF3-FLAG expression construct at 24 hours post-infection. Cells were then fixed at 48 hours post-infection for indirect immunofluorescence staining of NS1 and FLAG followed by confocal imaging. FLAG-tagged TM9SF3 displayed juxtanuclear and cytoplasmic staining profiles consistent with Golgi-associated localization (Fig. 5G). The co- localization between TM9SF3 and either the intense juxtanuclear NS1 foci or punctate cytoplasmic NS1 was infrequent, with line profile analysis showing only occasional overlap between TM9SF3- and NS1-associated peaks (Fig. 5H and 5I). The limited co-localization observed between TM9SF3 and NS1 suggests that these proteins may only transiently interact or be in close proximity for a small subset of the intracellular NS1 population. These findings also support the possibility that TM9SF3 is required for DENV infectious virion production and intracellular NS1 abundance and secretion through indirect mechanisms.

**Figure 5.**
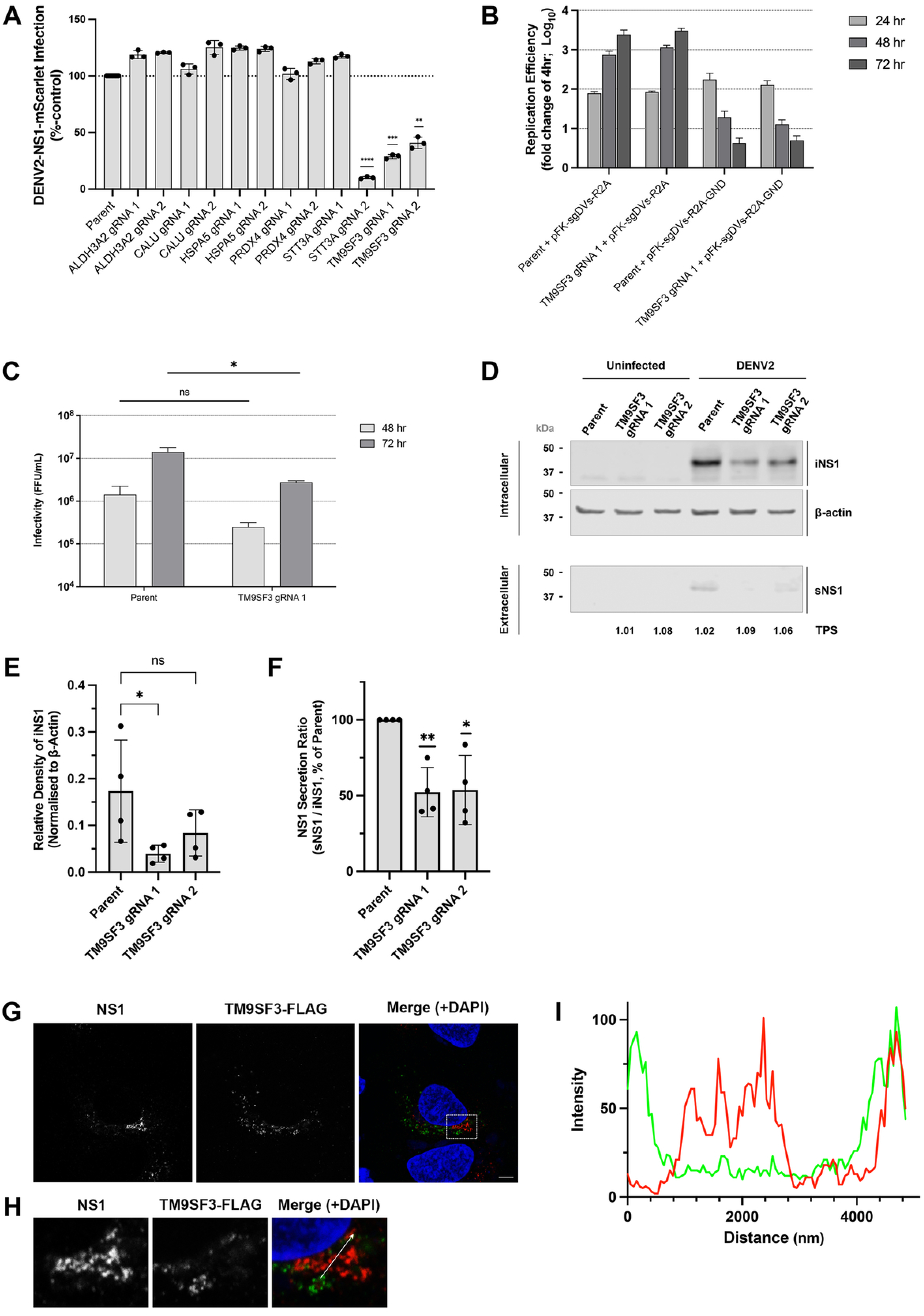
TM9SF3 knockout reduces DENV infection levels, infectious virus production and intracellular NS1 abundance and secretion. (A) Parent Huh-7.5 cells or CRISPR-Cas9-treated Huh-7.5 cells targeting proteins identified as hits in the siRNA screen were infected with DENV2-NS1-mScarlet (MOI: ∼0.17) and imaged every 3 hours over a 48-hour period via automated live-cell imaging. Mean mScarlet fluorescence area (µm2/image) was quantified and normalised to that of the parent control. Data are means ± S.D. (n=3). One sample *t*-test (**p < 0.005, ***p < 0.001, ****p < 0.0001). (B) Replication efficiency in parent or TM9SF3 knockout Huh-7.5 cells transfected with sgDVS-R2A or sgDVS-NS1-APEX2-R2A RNA. Cells were transfected with the indicated IVT DENV2 subgenomic replicon RNA transcripts and lysates were harvested at 4, 24, 48 and 72 hours post-transfection for measurement of *Renilla* luciferase activity. Raw luciferase values were normalised to the 4-hour timepoint average value (representative of input RNA). Replication deficient mutants harbouring a ‘GND’ mutation within NS5 were included as negative controls. Data are means ± S.D. (n=3). (C) Infectious virus production in parent or TM9SF3 knockout Huh-7.5 cells. Cells were infected with DENV2 at an MOI of 0.1, washed and returned to culture prior to the collection of supernatants at 48 and 72 hours post-infection. Infectivity of virus within cell supernatants was determined by FFA. Data are means ± S.D. (n=3). Unpaired *t*-test (*p < 0.05). (D) Parent or TM9SF3 knockout Huh-7.5 cells were infected with DENV at an MOI of 0.1. At 48 hours post-infection, cell culture supernatants and lysates were recovered to measure secreted NS1 (sNS1) and intracellular NS1 (iNS1) levels. Total proteins from supernatants and lysates were separated by SDS-PAGE and analysed by Western blotting using anti-NS1 and anti-β-actin. (E) Quantification of NS1 abundance in cell lysates by quantitative Western blot analysis, displayed as relative density of iNS1 normalised to that of β-actin. Data are means ± S.D. (n=4). Kruskal-Wallis test (*p < 0.05). (F) Quantification of NS1 abundance in cell supernatants by quantitative Western blot analysis, displayed as the secretion ratio of NS1 (sNS1 / iNS1) as a percentage of that of the parent control. Data are means ± S.D. (n=4). One sample *t*-test (*p < 0.05, **p < 0.01). (G) Localization of NS1 with respect to TM9SF3-FLAG. Huh-7.5 cells were infected with DENV2 (MOI: ∼0.1) and transfected with a TM9SF3-FLAG expression construct at 24 hours post-infection. At 48 hours post-infection, cells were fixed and subjected to indirect immunofluorescent labelling using mouse anti-NS1, rabbit anti-FLAG, Alexa Fluor 555-conjugated anti-mouse IgG (red) and Alexa Fluor 488-conjugated anti-rabbit IgG (green). Samples were stained with DAPI (blue) and analyzed by confocal fluorescence microscopy. Scale bar represents 5 µm. (H) Insets showing zoomed-in images of the white boxed region in the merged image shown in G. (I) Line profile corresponding to the 15 µm white arrow in the inset of the merged image in H.

## DISCUSSION

The identification of interactions between DENV NS1 and host proteins is essential for improving our understanding of the many critical functions of NS1 and may uncover targets for future antiviral therapies. Previous immunoprecipitation- and affinity purification-coupled proteomics studies have provided valuable insights into NS1-host factor interactions and NS1 biology (30, 42–44). However, given the multifunctional nature of NS1, it is likely that many interactions, including transient and weak interactions, remain unidentified. Furthermore, these studies primarily relied on subgenomic replicons or viral protein overexpression and therefore were unable to capture interactions during a productive viral infection. There remains a need for the development of approaches that enable the mapping of virus-host interactions throughout the complete viral replication cycle. Here, we used infectious APEX2-tagged DENV reporter viruses in combination with APEX2 proximity labelling-coupled quantitative proteomics to map the proteomic composition of the NS1 microenvironment in live infected Huh-7.5 hepatoma cells. Unlike immunoprecipitation- and affinity purification-based approaches, APEX2 proximity labelling has the potential to capture both stable and transient NS1-host factor interactions, as well as proteins that are closely proximal to but do not directly interact with NS1. The extent of protein biotinylation is dependent upon proximity to APEX2 and the availability of accessible electron-rich amino acid residues with which biotin-phenoxyl radicals react (45). Consequently, proteins that are closely proximal to and/or directly interact with NS1-APEX2 are expected to undergo greater biotinylation and therefore exhibit greater enrichment than proximal non-interacting proteins.

Our quantitative proteomic analysis resulted in the identification of 51 NS1-proximal host proteins, including 14 proteins that have previously been reported as DENV HDFs and 3 proteins previously identified as being critical to aspects of NS1 biology. A substantial proportion of the identified NS1-proximal proteins reside within the ER lumen or associate with ER membranes, consistent with the predominant localization of intracellular NS1 dimers to virus-induced ER-derived viral replication organelles (9, 10). Gene Ontology analysis further demonstrated that many of the identified NS1-proximal host proteins function in cellular pathways that have been widely implicated in DENV infection, including protein folding, post-translational protein modification and lipid metabolism and cholesterol homeostasis (3). Apart from 7 proteins (AADAC, AFP, ALDH3A2, AMBP, LIPC, PCSK9 and PROS1) that are predominantly expressed by hepatocytes, the identified NS1-proximal host proteins are broadly expressed across cell types targeted by DENV. Together, these results demonstrate the capacity of the utilised APEX2 proximity labelling-coupled quantitative proteomics approach described here to capture biologically relevant NS1-associated host proteins.

An siRNA screen was performed to determine whether select NS1-proximal host proteins are required for efficient DENV infection. The top hits of this screen included the well-characterized orthoflaviviral HDFs HSPA5 and STT3A. Numerous studies have demonstrated that the depletion or proteolytic cleavage of HSPA5 (also known as BiP or GRP78), a member of the HSP70 family, substantially reduces viral protein abundance and infectious virion production for DENV and other orthoflaviruses Zika virus (ZIKV) and Japanese encephalitis virus (JEV) (32, 46–55). The siRNA-mediated depletion of HSPA5 or pharmacological inhibition of its upstream transcriptional regulators was also shown to impair DENV NS1 folding and secretion (32, 56). Furthermore, interactions between HSPA5 and NS1 or E have been reported for DENV, ZIKV and JEV (32, 46, 50, 52, 53, 55, 57). STT3A is one of the two catalytic subunit isoforms of the OST complex, which catalyses *N-* linked glycosylation. OST complexes containing STT3A mediate the co-translational *N*- glycosylation of glycoproteins, whereas those containing STT3B modify *N*-glycosylation sites not processed by STT3A, acting to complete glycosylation co- or post-translationally (39, 58). Numerous CRISPR-Cas9 and siRNA screens have demonstrated that the depletion of STT3A strongly inhibits DENV infection (30, 39–41). The combined depletion or pharmacological inhibition of OST catalytic subunits STT3A and STT3B has been shown to substantially impair the *N*-glycosylation of NS1 and DENV replication (30). Other studies have indicated that STT3A and STT3B support DENV infection through glycosylation-independent mechanisms. These non-canonical functions are proposed to involve the stabilisation of the viral replication complex or other host proteins required for efficient viral replication (39, 41, 59). In addition to HDFs HSPA5 and STT3A, 4 host proteins not previously shown to be involved in DENV infection were identified as hits in our siRNA screen, including the aldehyde dehydrogenase ALDH3A2, calcium-binding chaperone CALU, peroxiredoxin PRDX4 and Golgi-resident transmembrane protein TM9SF3. A significant increase in DENV infection levels was observed for the siRNA-mediated knockdown of host proteins not previously characterized as restriction factors, possibly due to the disruption of ER homeostasis and stress-induced responses that DENV can exploit to enhance viral replication (3).

We further investigated the role of TM9SF3, a transmembrane receptor that mediates the selective autophagy of Golgi fragments in response to various forms of stress to preserve Golgi and glycosylation homeostasis (34). To achieve this, TM9SF3 binds through its N-terminal LC3-interacting regions (LIRs) to ATG8-family proteins, which targets Golgi fragments for engulfment by phagophores and subsequent lysosomal degradation (34). The knockout of TM9SF3, the mutation of its LIRs, or the knockout of the ATG8-activating enzyme ATG7 prevented Golgi fragmentation and targeting to lysosomes and resulted in the intracellular accumulation of incompletely glycosylated proteins. Conversely, TM9SF3 overexpression has been shown to increase the degradation of incompletely glycosylated proteins (34). TM9SF3 knockout has also been shown to reduce co-localization between the cis-Golgi and trans-Golgi network (TGN) and impair the maturation of high-mannose glycans into complex glycan forms (33). Considering the role of TM9SF3 in maintaining Golgi and protein glycosylation homeostasis, it is possible that the depletion of this protein disrupts Golgi-dependent stages of the DENV lifecycle. This could include the maturation and release of virions, the glycosylation of DENV glycoproteins E, NS1 and NS4B and the glycosylation and trafficking of HDFs. Given that TM9SF3 was confidently identified as NS1- proximal, it may be an important determinant of NS1 maturation, glycosylation and secretion; processes which are strongly associated with the Golgi (60). Using *Renilla* luciferase-encoding subgenomic DENV replicons, we first demonstrated that TM9SF3 knockout does not impact viral RNA replication. However, measurement of viral infectivity revealed a reduction in infectious virion production in TM9SF3 knockout cells. These findings suggest that the observed reduction in DENV infection following TM9SF3 knockdown in our siRNA screen may be attributable to defects in infectious virion production and spread. Furthermore, we demonstrated by quantitative Western blot analysis that TM9SF3 knockout reduced intracellular NS1 abundance and secretion. This could arguably be attributable to TM9SF3 knockout-associated incomplete or aberrant *N*-glycosylation of NS1 and subsequent effects on its folding, oligomerization and/or stability and/or the reduced detection of mature, *N*-glycosylated NS1 using the conformation-specific NS1-targeting monoclonal antibody employed here. Extension of our findings using additional loss-of-function approaches and further mechanistic analysis is required to clarify whether TM9SF3 is directly hijacked and manipulated by DENV NS1 or whether TM9SF3 depletion indirectly impacts upon NS1 maturation and secretion and infectious virus production. Further investigations of the impact of TM9SF3 on NS1 glycosylation, and whether a direct interaction occurs between these proteins, is also warranted. Furthermore, given the ubiquitous expression of human TM9SF3 and the expression of orthologs in DENV mosquito vectors, the role of this protein in the DENV lifecycle should be assessed across additional cell types, as well as in the context of infection with additional DENV serotypes and related orthoflaviviruses.

In summary, this study has demonstrated the utility of applying infectious APEX2- tagged reporter viruses and APEX2 proximity labelling-coupled quantitative proteomics to the identification of novel host proteins, including TM9SF3, with functional relevance to the DENV lifecycle and NS1 biology. Such virus-host interactions may be considered as future targets for the development of antiviral therapies.

## MATERIALS AND METHODS

### Cell Culture

Huh-7.5 cells were generously provided by Charles M. Rice (Rockefeller University, New York, USA) and were maintained as described previously (61, 62).

### Antibodies and Chemicals

Mouse anti-NS1 monoclonal antibody (MAb) 4G4 was generously provided by Jody Peters and Roy Hall (University of Queensland, Brisbane, Australia) or purchased from Mozzy Mabs (University of Queensland, Brisbane, Australia) (63). The mouse anti-Envelope monoclonal antibody was prepared from the hybridoma cell line D1-4G2-4-15 (4G2) purchased from ATCC and maintained as described previously (64). Mouse anti-β-actin MAb (AC-15) was purchased from Sigma-Aldrich. Rabbit anti-FLAG (D4W5B) was purchased from Cell Signalling Technologies. Alexa Fluor 488- and 555-conjugated anti-mouse IgG and anti-rabbit IgG secondary antibodies were purchased from Thermo Fisher Scientific. Cy3-conjugated streptavidin was purchased from Jackson ImmunoResearch Labs. Fluorescent stain 4’,6-diamidino-2-phenylindole (DAPI) was purchased from Sigma- Aldrich. Revert™ 700 Total Protein Stain, IRDye® 800 CW goat anti-mouse secondary antibody, and IRDye® 800 CW streptavidin for Western Immunoblotting were purchased from LI-COR Biosciences.

### Virus Plasmids and Virus Propagation

Plasmid pFK-DVs, containing a full-length DENV2 genome (strain 16681), the derivative full-length reporter construct pFK-DVs-R2A encoding a Renilla luciferase reporter gene (R2A), the subgenomic replicon derivative pFK-sgDVs-R2A encoding R2A, and the pFK-sgDVs- R2A replication-deficient NS5 mutant derivative pFK-sgDVs-GND-R2A were generously provided by Ralf Bartenschlager (University of Heidelberg, Heidelberg, Germany) (65). Reporter viruses pFK-DVs-NS1-APEX2 and pFK-DVs-NS1-mScarlet, bearing APEX2 and mScarlet tags within NS1, respectively, have been described previously (10). To generate a DENV2 reporter virus encoding a self-cleaved APEX2 tag (pFK-DVs-2A-APEX2-2A), a synthetic gene fragment (gBlock: Integrated DNA Technologies) encoding the APEX2 tag flanked by a 5’ P2A and 3’ T2A self-cleaving peptide sequence was inserted into the pFK-DVs-R2A backbone, which had been linearized with *Asc*I and *Age*I, using NEBuilder HiFi DNA Assembly master mix (New England Biolabs). The nucleotide sequence was confirmed by Sanger sequencing (AGRF). To initiate viral replication and produce infectious DENV2 stocks, RNA was transcribed *in vitro* from *Xba*I-linearized DENV2 plasmids using the mMessage mMachine SP6 transcription kit (Thermo Fisher Scientific) and transfected into Huh-7.5 cells using the DMRIE-C reagent (Thermo Fisher Scientific) as described previously (62, 64). Virus infectivity was measured by focus-forming assays as described previously, with the modification that an anti-Envelope MAb was used for immunofluorescent labelling (10).

### APEX2 Proximity Labelling

SILAC and APEX2 proximity labelling was performed as described previously (28). SILAC media (Thermo Fisher Scientific) deficient in L-lysine and L-arginine was supplemented with 10% (v/v) fetal calf serum (FCS; Sigma-Aldrich), penicillin–streptomycin–glutamine (Gibco, Invitrogen), 4.5 g/L D-(+)-glucose (Sigma-Aldrich) and either heavy, medium, or light isotopes of L-lysine (146 mg/L) and L-arginine (84 mg/L). Three Huh-7.5 cell cultures were maintained in heavy-, medium- and light-isotope SILAC media for 7 cell doublings. Cells were then transfected with IVT DENV2-NS1-APEX2 or DENV2-2A-APEX2-2A RNA using the DMRIE-C reagent (Thermo Fisher Scientific) alongside a mock-transfected negative control. In the first experiment, heavy-, medium- and light isotope-labelled cells were transfected with DENV2-NS1-APEX2 IVT RNA or DENV2-2A-APEX2-2A IVT RNA, or mock-transfected, respectively. SILAC label-switching was performed in subsequent replicates (Fig. 2B). To monitor infection levels, aliquots from each culture were sub-cultured at the 4- day post-transfection passage and were fixed, stained for viral E protein using the mouse anti-Envelope MAb (4G2) and Alexa Fluor 488-conjugated anti-mouse IgG secondary antibody, and counter-stained with DAPI. Infection levels were quantified as the proportion of E-positive cells relative to the total number of nuclei using a Cytation 5 cell imaging multi- mode reader (BioTek). At 5 days post-transfection, cells were incubated in SILAC media containing with 500 μM biotin-phenol (Sigma-Aldrich) for 30 minutes at 37°C. To initiate biotinylation, H₂O₂ (Sigma-Aldrich) was added to a final concentration of 1 mM and cells were incubated for 1 minute at room temperature. Reactions were quenched by 5 sequential 1-minute washes with Dulbecco’s PBS (DPBS; Sigma-Aldrich) supplemented with sodium ascorbate (10 mM final concentration; Sigma-Aldrich), sodium azide (10 mM final concentration; Sigma-Aldrich) and Trolox (5 mM final concentration; Sigma-Aldrich). Cells were then lysed in ice-cold RIPA buffer supplemented with protease inhibitors, sodium ascorbate (10 mM), sodium azide (10 mM) and Trolox (5 mM), homogenized, and clarified by centrifugation at 10,000 x g for 10 min at 4°C.

### Proteomic Analysis of Biotinylated Proteins

The concentration of proteins in lysates harvested from heavy, medium and light isotope-labelled cells was determined using the Pierce 660 nm assay (Thermo Fisher Scientific). To normalize these values, equal concentrations of each lysate were resolved by SDS-PAGE and total protein was quantified by Oriole fluorescent protein stain (Bio-Rad). Streptavidin pulldown was performed as described previously (28). Lysate samples were mixed in a 1:1:1 ratio by protein mass (2 mg of protein per sample) and incubated with pre-equilibrated Pierce streptavidin magnetic beads (Thermo Fisher Scientific) overnight at 4°C with gentle rotation. Beads were washed sequentially: twice with RIPA buffer, once with 1 M KCl, once with 0.1 M Na₂CO₃, once with 2 M urea in Tris-HCl (pH 8.0), and twice in RIPA buffer. Bound proteins were eluted by boiling in 3X SDS-PAGE loading buffer containing 2 mM biotin at 95°C for 10 min, separated by SDS-Page using NuPAGE Novex 4–12% Bis-Tris gels (Thermo Fisher Scientific) and visualized using SYPRO Ruby protein stain (Bio-Rad). Each lane was cut into 22 pieces for in-gel tryptic digestion. Gel pieces were washed in water, destained with 1:1 (v/v) acetonitrile (ACN)/50 mM ammonium bicarbonate (ABC) and dehydrated in 100% ACN. Proteins were reduced with 50 mM dithiothreitol (DTT) in 50 mM ABC at 56°C for 20 minutes and alkylated with 100 mM iodoacetamide (IAA) in 50 mM ABC in the dark at room temperature for 20 minutes. Gel pieces were then washed in water, dehydrated sequentially in 1:1 (v/v) ACN/50 mM ABC and dried at 37°C for 10 minutes. Peptides were digested with trypsin (12.5 ng/µL; Thermo Fisher Scientific) in 50 mM ABC/50 mM CaCl₂ overnight at 37°C. Peptides were collected, acidified with 0.05% formic acid, dried by vacuum centrifugation, and stored at –20°C. Prior to LC-MS/MS, peptides were resuspended in 5% ACN/0.1% formic acid in mass spectrometry-grade water and transferred to glass screw-cap vials (Thermo Fisher Scientific).

LC-MS/MS was performed using an Orbitrap Exploris 480 mass spectrometer coupled to a Dionex UltiMate 3000 RSLCnano uHPLC system and Dionex UltiMate 3000 RS Autosampler (Thermo Fisher Scientific). Samples were loaded onto an in-house packed pre-column (150 µm x 10 mm, ReproSil-Pur 120 C18-AQ, 3 µm beads) for desalting and then separated on an in-house packed analytical column (75 µm × 25 cm, ReproSil-Pur 120 C18-AQ, 3 µm beads) at 300 nL/min using a 140-min gradient (3–33% buffer B over 120 min, ramp to 99% buffer B over 5 min; buffer A: 0.1% formic acid in water; buffer B: 0.1% formic acid in acetonitrile). The mass spectrometer was operated in data-dependent acquisition mode with a 3 second cycle time. MS1 scans were acquired at a resolution of 120,000 (m/z 350–1400, AGC 2x10⁶). Top precursors (charge states of 2-7) were isolated (1.4 m/z) and fragmented by HCD (30% normalized collision energy). MS2 scans were acquired at a resolution of 15,000 (AGC 1x10⁵, max injection 200 ms) with dynamic exclusion enabled (75 second duration, ±10 ppm tolerance). Data were processed using Proteome Discoverer (version 2.4.1.15). Raw data was searched using Sequest HT against a custom database containing the Swiss-Prot human proteome, DENV2 and DENV2-NS1-APEX2 protein sequences, and common contaminants. Search parameters were set as follows: trypsin digestion allowing up to 2 missed cleavages; static modification: carbamidomethylation (Cys, +57.021 Da); dynamic modifications: methionine oxidation (+15.995 Da), Asn/Gln deamidation (+0.984 Da), Glu methylation (+14.016 Da), BP labelling (His, Trp, Tyr, +361.146 Da), and desthiobiotin-phenol labelling (Tyr, +331.190 Da). Peptides were filtered using Percolator node with a false discovery rate (FDR) threshold of 1% and quantified using the Minora Feature Detector with SILAC 3-plex labelling (medium: Lys4, Arg6; heavy: Lys8, Arg10).

### Bioinformatics, Statistical Analysis and Figure Preparation

Data were processed in the R statistical programming environment using Tidyverse (version 2.0.0) (66, 67). Log₂-transformed raw abundances were normalized with HarmonizR (version 1.2.0) to correct for batch effects between runs and systematic differences between SILAC labelling conditions (68). Samples were visualized using classical multidimensional scaling (MDS) before and after normalization. Log₂ raw abundances for all replicate pairs were also plotted as scatter plots to assess normalization. Log₂ fold changes were calculated between DENV2-NS1-APEX2, DENV2-2A-APEX2-2A, and mock conditions using the limma package, and statistical significance was determined with t-tests using rstatix (version 0.7.2) with Holm–Bonferroni p-value adjustment (69, 70). Final volcano plots and heatmaps were generated with ggplot2 (version 3.5.1) (71). Gene Ontology analysis was performed using the fgsea package (version 1.26.0) (72). Unless otherwise indicated, GraphPad Prism (version 10.2.2) was employed for statistical analysis and graphing and figures were compiled using Adobe Photoshop (version 24.1.0).

### siRNA Screening

A custom cherry-pick ON-TARGETplus siRNA library (Dharmacon, Horizon Discovery) comprising 40 SMARTpools was used. Each SMARTpool contained four siRNAs targeting distinct sequences within a single gene. A non-targeting scrambled siRNA (NTC) was included as a negative control, and a custom siRNA duplex targeting the DENV2 NS5 gene served as a positive control for inhibition of viral replication. The siRNAs were resuspended to 1 µM in 1X siRNA buffer (Dharmacon, Horizon Discovery), and 4 µL (4 pg) of each SMARTpool or control was dispensed into 96-well assay plates (PhenoPlates, PerkinElmer) using a Janus G3 automated liquid handler (PerkinElmer). Plates were sealed and stored at –80°C until use. Three independent siRNA screens were performed, each consisting of two siRNA assay plates. On each plate, SMARTpools were tested in duplicate, and both negative and positive controls were each included in eight wells per plate.

Huh-7.5 cells were reverse-transfected with the siRNA library. DharmaFECT 4 (Dharmacon, Horizon Discovery) was diluted in Opti-MEM to prepare a transfection master mix. The transfection mix was added to the siRNA-containing assay plates (0.6 µL of DharmaFECT 4 in 19.4 µL of Opti-MEM per well, 16 µL per well total) and incubated at room temperature for 20 minutes with agitation to allow siRNA-DharmaFECT complex formation. Huh-7.5 cells were trypsinized, resuspended in antibiotic-free culture media and added to the assay plates at a density of 1.25 x 104 cells per well in 80 µL, yielding a final siRNA concentration of 40 nM per well. Cells were cultured for 3 hours at 37°C, after which the media containing siRNAs was replaced with complete media. At 20 hours post-transfection, cells were infected with DENV2-NS1-mScarlet virus (MOI ∼0.25) and cultured for 3 hours at 37°C, after which the inoculum was removed and replaced with complete media. At 48 hours post-infection, cells were fixed in ice-cold acetone:methanol (1:1) for 5 minutes at 4°C, washed in 1X PBS, counter-stained with DAPI (1 µg/mL) for 10 minutes in the dark, and stored in 1X PBS at 4°C. Cells were imaged using an Operetta CLS high-content imaging system (PerkinElmer) controlled by Harmony software (version 5.3; Revvity). Images were segmented and nuclei counts and mScarlet fluorescence intensity were quantified using the Columbus (version 2.5; Revvity) image data storage and analysis system running on Acapella (version 4.0) and OMERO servers (version 4.4.1.1, Revvity).

### Generation of Knockout Cell Lines

Pre-designed 20-nucleotide gRNA sequences were selected using the IDT Predesigned Alt-R CRISPR-Cas9 guide RNA design web tool. The gRNAs were synthesized as single-stranded DNA oligonucleotides and annealed prior to ligation into the *Bsm*BI-v2-digested LentiCRISPRv2 plasmid. The plasmids were then packaged into lentiviruses for the transduction of Huh-7.5 cells as described previously (73). Transduced cells were selected using puromycin dihydrochloride (3 µg/mL; Sigma-Aldrich) for three weeks. The surviving polyclonal cell lines were then used in live-cell imaging-based screens, luciferase assays, infectivity assays and Western blotting experiments, as indicated.

### Infectivity Assays

Cell culture supernatants containing DENV were collected at the indicated time points and stored at –80°C. Infectious virus was titrated by focus forming assay, as described previously (10).

### Quantification of Subgenomic DENV RNA Replication by Luciferase Assay

Huh-7.5 cells or Huh-7.5 TM9SF3 knockout cells were seeded into 12-well plates at 6.5 x 104 cells per well and cultured overnight. The cells were then transfected with IVT *Renilla*- luciferase encoding subgenomic DENV2 replicon RNA using DMRIE-C (Thermo Fisher Scientific) according to the manufacturer’s instructions. At 4-, 24-, 48-, 72- and 96-hours post-transfection, the cell culture media was removed, and the monolayers were washed once with 1X PBS, lysed in 1X Renilla Luciferase Assay Buffer (Promega) and stored at - 20°C. Following thawing at room temperature, lysates were transferred to a 96-well white OptiPlate (PerkinElmer). *Renilla* luciferase activity was then measured using a Cytation 5 cell imaging multi-mode plate reader equipped with a Dual-Reagent Injector Module (BioTek) following the automated addition (2 s delay, 10 s integration) of 1X Renilla Luciferase Assay Substrate (Promega).

### SDS-PAGE and Western Blotting

Cell culture supernatants were diluted 1:1 in ice-cold RIPA lysis buffer containing 1X protease inhibitor cocktail (Sigma-Aldrich). Cell monolayers were washed once with 1X PBS and lysed in RIPA lysis buffer supplemented with 1X protease inhibitor cocktail for 20 minutes on ice. Lysates were homogenized using a needle, clarified by centrifugation (10,000 x g, 10 minutes, 4°C) and mixed with either reducing or non-reducing 1X SDS-PAGE loading buffer. Proteins were denatured at 95°C for 5 minutes, separated by SDS-PAGE and transferred to nitrocellulose membranes (Bio-Rad). For cell culture supernatant samples, total protein normalisation was then performed using Revert 700 Total Protein Stain (LI-COR Biosciences) according to the manufacturer’s instructions and membranes were imaged using an Odyssey Infrared Imaging System (LI-COR Biosciences). Membranes were blocked in 5% (w/v) skim milk in Tris-buffered saline (TBS) for 1 hour at room temperature and then incubated overnight at 4°C with primary anti-NS1 (MAb 4G4; 1:5 dilution) and anti-β-actin (MAb AC-15; 1:10,000) antibodies diluted in 1% (w/v) skim milk in TBS-T (TBS containing 0.1% [v/v] Tween-20). Following washes in TBS-T, the membranes were incubated with IRDye® 800 CW goat anti-mouse IgG and 680RD goat anti-rabbit IgG secondary antibodies (1:20,000) or IRDye® 800CW streptavidin (1:1000) diluted in TBS-T. Following washes in TBS-T, the membranes were imaged using an Odyssey Infrared Imaging System (LI-COR Biosciences). Protein signal intensities were quantified using Image Studio Lite (version 5.2.5).

### Generation of the TM9SF3-FLAG Expression Construct

To generate the TM9SF3-FLAG expression construct, the pLenti6/V5-D-Topo vector (Thermo Fisher Scientific) was first linearized using *Bam*HI and *Mlu*I restriction enzymes (New England Biolabs). A gene fragment encoding TM9SF3 (NM_020123.4) fused to the FLAG epitope via a flexible Glycine-Serine-Glycine linker and containing 20-nucleotide 5’ and 3’ homology arms matching the linearised vector, was synthesized (Integrated DNA Technologies). The NEBuilder HiFi DNA Assembly Master Mix (New England Biolabs) was used to assemble the TM9SF3-FLAG gene fragment into the linearized pLenti6/V5-D-Topo vector. Following transformation and plasmid DNA propagation, the cloned cDNA sequence was verified by Sanger Sequencing (Australian Genome Research Facility).

### Immunofluorescent Labelling and Confocal Microscopy

Huh-7.5 cells were seeded at 1 x 104 cells per well into 0.2% (w/v) gelatine-coated 8-well #1.5H coverslip-bottom chambered coverslips (μ-Slide, Ibidi Gmbh) and returned to culture overnight. For analysis of NS1 and biotinylated protein co-localization, the cells were transfected with DENV2-2A-APEX2-2A or DENV2-NS1-APEX2 IVT RNA using DMRIE-C (Thermo Fisher Scientific) according to the manufacturer’s instructions and then treated with BP/H2O2 at 5 days post-transfection. For analysis of NS1 and TM9SF3 co-localization, cells were infected with DENV (MOI: ∼0.1) and then transfected with the TM9SF3-FLAG expression construct at 24 hours post-infection using Lipofectamine 3000 (Invitrogen) according to the manufacturer’s instructions. To perform indirect immunofluorescent labelling, cell monolayers were washed with 1X PBS and then fixed with ice-cold acetone methanol (1:1) for 5 minutes on ice. The fixative was removed and the samples were washed once with 1X PBS before blocking in 5% (w/v) BSA (Sigma-Aldrich) in PBS for 30 minutes at room temperature. Following blocking, the samples were incubated overnight at 4°C with the indicated primary antibodies (anti-FLAG [MAb D4W5B, 1:200] and/or anti-NS1 [MAb 4G4; 1:5]) diluted in 1% BSA in 1X PBS. The samples were washed with 1X PBS and then incubated for 1 hour at 4°C in the dark with the indicated secondary antibodies (Alexa Fluor 488-conjugated anti-mouse IgG and streptavidin-conjugated Cy3 or Alexa Fluor 555- conjugated anti-mouse IgG and Alexa Fluor 488-conjugated anti-rabbit IgG; Thermo Fisher Scientific) diluted 1:100-1:200 in 1% BSA in 1X PBS. Following washing in 1X PBS, the samples were counterstained with DAPI (Sigma-Aldrich) diluted to 1 µg/mL in 1X PBS for 10 minutes at room temperature in the dark. The stain was replaced with 1X PBS, and the samples were imaged using a ZEISS LSM 880 Fast Airyscan Confocal Microscope with a C-Plan-Apochromatic 63× (NA: 1.4) oil immersion objective lens. For this, 405, 488 and 561 nm laser lines were used at 2% of maximum power, with master gain settings adjusted to enable visualisation of signals with minimal saturation and pinhole sizes adjusted to 1.0 Airy units for the longest-wavelength fluorophore and matched for all tracks. Images were processed and analyzed using the ZEISS ZEN Blue software (version 3.2).

## ACKNOWLEDGEMENTS

We thank the following people for generously providing reagents (as detailed in Materials and Methods): Charles M. Rice (Rockefeller University); Ralf Bartenschlager (University of Heidelberg); Jody Peters (University of Queensland) and; Roy Hall (University of Queensland). We acknowledge and extend appreciation to the staff and for the instrumentation and services provided at the following research facilities that were used during this study: Flinders Microscopy and Microanalysis (Flinders University); Cell Screen SA Facility (Flinders University); Flinders Proteomics Facility (Flinders University); and the Australian Genome Research Facility (Adelaide, South Australia). We acknowledge the facilities and the scientific and technical assistance of Microscopy Australia (ROR: 042mm0k03), enabled by NCRIS and the government of South Australia at Flinders Microscopy and Microanalysis (ROR: 04z91ja70), Flinders University (ROR: 01kpzv902). We acknowledge funding support from the National Health and Medical Research Council (NHMRC, Australia) (2012370) to N.S.E.

---

**Supplementary Figure 1. Confirmation of APEX2-mediated protein biotinylation by streptavidin blotting prior to proteomic analysis.** Huh-7.5 cells maintained in heavy, medium or light SILAC media were either transfected with *in vitro* transcribed DENV2-NS1- APEX2 or DENV2-2A-APEX2-2A RNA, or mock-transfected. At 5 days post-transfection, the cells were treated with biotin-phenol and H2O2 prior to lysis. Three independent experiments were performed. Total proteins from lysed cells were separated by SDS-PAGE and analysed by Western blotting using fluorescently labelled streptavidin, anti-NS1 and anti-β-actin.

**Supplementary Figure 2. Fractionation of biotinylated proteins isolated by streptavidin pulldown.** Biotinylated proteins from three independent APEX2 proximity labelling experiments were enriched using streptavidin-coated magnetic beads and eluted by boiling in SDS-PAGE sample buffer supplemented with biotin. For each replicate, eluates were loaded across two adjacent lanes on an SDS-PAGE gel. Following separation, proteins were visualised by SYPRO ruby staining, and 22 bands were excised from each lane for in- gel tryptic digestion of proteins and LC-MS/MS analysis.

**Supplementary Figure 3. Pairwise correlation of normalised protein abundances across independent biological replicates.** Scatter plots illustrating pairwise comparisons of log₂-transformed, normalized protein abundances between the three independent experimental replicates for each experimental condition. Row 1: DENV2-NS1-APEX2 replicate comparisons; Row 2: DENV2-2A-APEX2-2A replicate comparisons; Row 3: mock replicate comparisons. Correlation coefficients (R²) for each replicate pair are indicated above each scatter plot.

**Supplementary Figure 4. Proteomic analysis of biotinylated NS1-proximal proteins.** Volcano plot of the relative enrichment of proteins in DENV2-NS1-APEX2 samples compared with mock samples. Proteins with log2 fold change > 1 and adjusted p-value < 0.05 are considered significantly enriched in DENV2-NS1-APEX2 samples and are highlighted in green (host proteins) or red (DENV2 proteins). Plot represents a merged dataset from three independent experiments.

**Supplementary Figure 5. Representative images of DENV2-NS1-mScarlet infection following siRNA treatment.** Images show a significant reduction in infection following the siRNA-mediated knockdown of ALDH3A2, CALU, HSPA5, PRDX4, STT3A and TM9SF3 in comparison to the NTC.

## REFERENCES

1. Bhatt S, Gething PW, Brady OJ, Messina JP, Farlow AW, Moyes CL, Drake JM, Brownstein JS, Hoen AG, Sankoh O, Myers MF, George DB, Jaenisch T, Wint GRW, Simmons CP, Scott TW, Farrar JJ, Hay SI. 2013. The global distribution and burden of dengue. Nature 496:504–507.

2. Brady OJ, Gething PW, Bhatt S, Messina JP, Brownstein JS, Hoen AG, Moyes CL, Farlow AW, Scott TW, Hay SI. 2012. Refining the Global Spatial Limits of Dengue Virus Transmission by Evidence-Based Consensus. PLOS Neglected Tropical Diseases 6:e1760.

3. Neufeldt CJ, Cortese M, Acosta EG, Bartenschlager R. 2018. Rewiring cellular networks by members of the Flaviviridae family. Nature Reviews Microbiology 16:125–142.

4. Winkler G, Maxwell SE, Ruemmler C, Stollar V. 1989. Newly synthesized dengue-2 virus nonstructural protein NS1 is a soluble protein but becomes partially hydrophobic and membrane-associated after dimerization. Virology 171:302–305.

5. Putnak JR, Charles PC, Padmanabhan R, Irie K, Hoke CH, Burke DS. 1988. Functional and antigenic domains of the dengue-2 virus nonstructural glycoprotein NS-1. Virology 163:93–103.

6. Flamand M, Megret F, Mathieu M, Lepault J, Rey Félix A, Deubel V. 1999. Dengue Virus Type 1 Nonstructural Glycoprotein NS1 Is Secreted from Mammalian Cells as a Soluble Hexamer in a Glycosylation-Dependent Fashion. Journal of Virology 73:6104–6110.

7. Pryor MJ, Wright PJ. 1994. Glycosylation mutants of dengue virus NS1 protein. J Gen Virol 75 ( Pt 5):1183–7.

8. Somnuke P, Hauhart RE, Atkinson JP, Diamond MS, Avirutnan P. 2011. N-linked glycosylation of dengue virus NS1 protein modulates secretion, cell-surface expression, hexamer stability, and interactions with human complement. Virology 413:253–64.

9. Mackenzie JM, Jones MK, Young PR. 1996. Immunolocalization of the Dengue Virus Nonstructural Glycoprotein NS1 Suggests a Role in Viral RNA Replication. Virology 220:232–240.

10. Eyre NS, Johnson SM, Eltahla AA, Aloi M, Aloia AL, McDevitt CA, Bull RA, Beard MR. 2017. Genome-Wide Mutagenesis of Dengue Virus Reveals Plasticity of the NS1 Protein and Enables Generation of Infectious Tagged Reporter Viruses. J Virol 91.

11. Gutsche I, Coulibaly F, Voss JE, Salmon J, d’Alayer J, Ermonval M, Larquet E, Charneau P, Krey T, Mégret F, Guittet E, Rey FA, Flamand M. 2011. Secreted dengue virus nonstructural protein NS1 is an atypical barrel-shaped high-density lipoprotein. Proceedings of the National Academy of Sciences 108:8003-8008.

12. Akey DL, Brown WC, Dutta S, Konwerski J, Jose J, Jurkiw TJ, DelProposto J, Ogata CM, Skiniotis G, Kuhn RJ, Smith JL. 2014. Flavivirus NS1 Structures Reveal Surfaces for Associations with Membranes and the Immune System. Science 343:881–885.

13. Shu B, Ooi JSG, Tan AWK, Ng T-S, Dejnirattisai W, Mongkolsapaya J, Fibriansah G, Shi J, Kostyuchenko VA, Screaton GR, Lok S-M. 2022. CryoEM structures of the multimeric secreted NS1, a major factor for dengue hemorrhagic fever. Nature Communications 13:6756.

14. Pan Q, Jiao H, Zhang W, Chen Q, Zhang G, Yu J, Zhao W, Hu H. 2024. The step-by-step assembly mechanism of secreted flavivirus NS1 tetramer and hexamer captured at atomic resolution. Science Advances 10:eadm8275.

15. Muller DA, Young PR. 2013. The flavivirus NS1 protein: molecular and structural biology, immunology, role in pathogenesis and application as a diagnostic biomarker. Antiviral Res 98:192–208.

16. Płaszczyca A, Scaturro P, Neufeldt CJ, Cortese M, Cerikan B, Ferla S, Brancale A, Pichlmair A, Bartenschlager R. 2019. A novel interaction between dengue virus nonstructural protein 1 and the NS4A-2K-4B precursor is required for viral RNA replication but not for formation of the membranous replication organelle. PLOS Pathogens 15:e1007736.

17. Scaturro P, Cortese M, Chatel-Chaix L, Fischl W, Bartenschlager R. 2015. Dengue Virus Non-structural Protein 1 Modulates Infectious Particle Production via Interaction with the Structural Proteins. PLOS Pathogens 11:e1005277.

18. Modhiran N, Watterson D, Muller DA, Panetta AK, Sester DP, Liu L, Hume DA, Stacey KJ, Young PR. 2015. Dengue virus NS1 protein activates cells via Toll-like receptor 4 and disrupts endothelial cell monolayer integrity. Science Translational Medicine 7:304ra142–304ra142.

19. Beatty PR, Puerta-Guardo H, Killingbeck SS, Glasner DR, Hopkins K, Harris E. 2015. Dengue virus NS1 triggers endothelial permeability and vascular leak that is prevented by NS1 vaccination. Sci Transl Med 7:304ra141.

20. Puerta-Guardo H, Glasner DR, Harris E. 2016. Dengue Virus NS1 Disrupts the Endothelial Glycocalyx, Leading to Hyperpermeability. PLoS Pathog 12:e1005738.

21. Puerta-Guardo H, Glasner DR, Espinosa DA, Biering SB, Patana M, Ratnasiri K, Wang C, Beatty PR, Harris E. 2019. Flavivirus NS1 Triggers Tissue-Specific Vascular Endothelial Dysfunction Reflecting Disease Tropism. Cell Reports 26:1598–1613.e8.

22. Puerta-Guardo H, Biering SB, de Sousa FTG, Shu J, Glasner DR, Li J, Blanc SF, Beatty PR, Harris E. 2022. Flavivirus NS1 Triggers Tissue-Specific Disassembly of Intercellular Junctions Leading to Barrier Dysfunction and Vascular Leak in a GSK- 3β-Dependent Manner. Pathogens 11.

23. Avirutnan P, Fuchs A, Hauhart RE, Somnuke P, Youn S, Diamond MS, Atkinson JP. 2010. Antagonism of the complement component C4 by flavivirus nonstructural protein NS1. J Exp Med 207:793–806.

24. Avirutnan P, Hauhart RE, Somnuke P, Blom AM, Diamond MS, Atkinson JP. 2011. Binding of Flavivirus Nonstructural Protein NS1 to C4b Binding Protein Modulates Complement Activation. The Journal of Immunology 187:424–433.

25. Thiemmeca S, Tamdet C, Punyadee N, Prommool T, Songjaeng A, Noisakran S, Puttikhunt C, Atkinson JP, Diamond MS, Ponlawat A, Avirutnan P. 2016. Secreted NS1 Protects Dengue Virus from Mannose-Binding Lectin-Mediated Neutralization. J Immunol 197:4053–4065.

26. Libraty DH, Young PR, Pickering D, Endy TP, Kalayanarooj S, Green S, Vaughn DW, Nisalak A, Ennis FA, Rothman AL. 2002. High Circulating Levels of the Dengue Virus Nonstructural Protein NS1 Early in Dengue Illness Correlate with the Development of Dengue Hemorrhagic Fever. The Journal of Infectious Diseases 186:1165–1168.

27. Lam SS, Martell JD, Kamer KJ, Deerinck TJ, Ellisman MH, Mootha VK, Ting AY. 2015. Directed evolution of APEX2 for electron microscopy and proximity labeling. Nat Methods 12:51–4.

28. Hung V, Udeshi ND, Lam SS, Loh KH, Cox KJ, Pedram K, Carr SA, Ting AY. 2016. Spatially resolved proteomic mapping in living cells with the engineered peroxidase APEX2. Nat Protoc 11:456–75.

29. Chapman-Smith A, Cronan JE, Jr. 1999. Molecular biology of biotin attachment to proteins. J Nutr 129:477s–484s.

30. Hafirassou ML, Meertens L, Umaña-Diaz C, Labeau A, Dejarnac O, Bonnet-Madin L, Kümmerer BM, Delaugerre C, Roingeard P, Vidalain PO, Amara A. 2017. A Global Interactome Map of the Dengue Virus NS1 Identifies Virus Restriction and Dependency Host Factors. Cell Rep 21:3900–3913.

31. Denolly S, Guo H, Martens M, Płaszczyca A, Scaturro P, Prasad V, Kongmanas K, Punyadee N, Songjaeng A, Mairiang D, Pichlmair A, Avirutnan P, Bartenschlager R. 2023. Dengue virus NS1 secretion is regulated via importin-subunit β1 controlling expression of the chaperone GRp78 and targeted by the clinical drug ivermectin. mBio 14:e0144123.

32. Songprakhon P, Limjindaporn T, Perng GC, Puttikhunt C, Thaingtamtanha T, Dechtawewat T, Saitornuang S, Uthaipibull C, Thongsima S, Yenchitsomanus PT, Malasit P, Noisakran S. 2018. Human glucose-regulated protein 78 modulates intracellular production and secretion of nonstructural protein 1 of dengue virus. J Gen Virol 99:1391–1406.

33. Tsui CK, Twells N, Durieux J, Doan E, Woo J, Khosrojerdi N, Brooks J, Kulepa A, Webster B, Mahal LK, Dillin A. 2024. CRISPR screens and lectin microarrays identify high mannose N-glycan regulators. Nature Communications 15:9970.

34. Yang J, Dong Y, Xu J, Qian X, Cai Y, Chen Y, Zhang P, Gao D, Cui Z, Cui Y. 2025. TM9SF3 is a Golgi-resident ATG8-binding protein essential for Golgi-selective autophagy. Dev Cell doi:10.1016/j.devcel.2025.06.017.

35. Kelson TL, Secor McVoy JR, Rizzo WB. 1997. Human liver fatty aldehyde dehydrogenase: microsomal localization, purification, and biochemical characterization. Biochim Biophys Acta 1335:99–110.

36. Fujii J, Ochi H, Yamada S. 2025. A comprehensive review of peroxiredoxin 4, a redox protein evolved in oxidative protein folding coupled with hydrogen peroxide detoxification. Free Radical Biology and Medicine 227:336–354.

37. Yabe D, Nakamura T, Kanazawa N, Tashiro K, Honjo T. 1997. Calumenin, a Ca2+- binding Protein Retained in the Endoplasmic Reticulum with a Novel Carboxyl-terminal Sequence, HDEF*. Journal of Biological Chemistry 272:18232–18239.

38. Wajih N, Sane DC, Hutson SM, Wallin R. 2004. The Inhibitory Effect of Calumenin on the Vitamin K-dependent γ-Carboxylation System: Characterisation of the System in Normal and Warfarin-Resistant Rats Journal of Biological Chemistry 279:25276–25283.

39. Marceau CD, Puschnik AS, Majzoub K, Ooi YS, Brewer SM, Fuchs G, Swaminathan K, Mata MA, Elias JE, Sarnow P, Carette JE. 2016. Genetic dissection of Flaviviridae host factors through genome-scale CRISPR screens. Nature 535:159–163.

40. Savidis G, McDougall WM, Meraner P, Perreira JM, Portmann JM, Trincucci G, John SP, Aker AM, Renzette N, Robbins DR, Guo Z, Green S, Kowalik TF, Brass AL. 2016. Identification of Zika Virus and Dengue Virus Dependency Factors using Functional Genomics. Cell Rep 16:232–246.

41. Lin DL, Cherepanova NA, Bozzacco L, MacDonald MR, Gilmore R, Tai AW. 2017. Dengue Virus Hijacks a Noncanonical Oxidoreductase Function of a Cellular Oligosaccharyltransferase Complex. mBio 8.

42. Cervantes-Salazar M, Angel-Ambrocio AH, Soto-Acosta R, Bautista-Carbajal P, Hurtado-Monzon AM, Alcaraz-Estrada SL, Ludert JE, Del Angel RM. 2015. Dengue virus NS1 protein interacts with the ribosomal protein RPL18: This interaction is required for viral translation and replication in Huh-7 cells. Virology 484:113–126.

43. Dechtawewat T, Paemanee A, Roytrakul S, Songprakhon P, Limjindaporn T, Yenchitsomanus PT, Saitornuang S, Puttikhunt C, Kasinrerk W, Malasit P, Noisakran S. 2016. Mass spectrometric analysis of host cell proteins interacting with dengue virus nonstructural protein 1 in dengue virus-infected HepG2 cells. Biochim Biophys Acta 1864:1270–1280.

44. Shah PS, Link N, Jang GM, Sharp PP, Zhu T, Swaney DL, Johnson JR, Von Dollen J, Ramage HR, Satkamp L, Newton B, Hüttenhain R, Petit MJ, Baum T, Everitt A, Laufman O, Tassetto M, Shales M, Stevenson E, Iglesias GN, Shokat L, Tripathi S, Balasubramaniam V, Webb LG, Aguirre S, Willsey AJ, Garcia-Sastre A, Pollard KS, Cherry S, Gamarnik AV, Marazzi I, Taunton J, Fernandez-Sesma A, Bellen HJ, Andino R, Krogan NJ. 2018. Comparative Flavivirus-Host Protein Interaction Mapping Reveals Mechanisms of Dengue and Zika Virus Pathogenesis. Cell 175:1931–1945.e18.

45. Qin W, Cho KF, Cavanagh PE, Ting AY. 2021. Deciphering molecular interactions by proximity labeling. Nature Methods 18:133–143.

46. Limjindaporn T, Wongwiwat W, Noisakran S, Srisawat C, Netsawang J, Puttikhunt C, Kasinrerk W, Avirutnan P, Thiemmeca S, Sriburi R, Sittisombut N, Malasit P, Yenchitsomanus PT. 2009. Interaction of dengue virus envelope protein with endoplasmic reticulum-resident chaperones facilitates dengue virus production. Biochem Biophys Res Commun 379:196–200.

47. Wati S, Soo ML, Zilm P, Li P, Paton AW, Burrell CJ, Beard M, Carr JM. 2009. Dengue Virus Infection Induces Upregulation of GRP78, Which Acts To Chaperone Viral Antigen Production. Journal of Virology 83:12871–12880.

48. Wu Y-P, Chang C-M, Hung C-Y, Tsai M-C, Schuyler SC, Wang RY-L. 2011. Japanese encephalitis virus co-opts the ER-stress response protein GRP78 for viral infectivity. Virology Journal 8:128.

49. Diwaker D, Mishra KP, Ganju L. 2015. Effect of modulation of unfolded protein response pathway on dengue virus infection. Acta Biochim Biophys Sin (Shanghai) 47:960–8.

50. Nain M, Mukherjee S, Karmakar SP, Paton AW, Paton JC, Abdin MZ, Basu A, Kalia M, Vrati S. 2017. GRP78 Is an Important Host Factor for Japanese Encephalitis Virus Entry and Replication in Mammalian Cells. J Virol 91.

51. Mukherjee S, Singh N, Sengupta N, Fatima M, Seth P, Mahadevan A, Shankar SK, Bhattacharyya A, Basu A. 2018. Japanese encephalitis virus induces human neural stem/progenitor cell death by elevating GRP78, PHB and hnRNPC through ER stress. Cell Death & Disease 8:e2556–e2556.

52. Chen J, Chen Z, Liu M, Qiu T, Feng D, Zhao C, Zhang S, Zhang X, Xu J. 2020. Placental Alkaline Phosphatase Promotes Zika Virus Replication by Stabilizing Viral Proteins through BIP. mBio 11:10.1128/mbio.01716-20.

53. Royle J, Ramírez-Santana C, Akpunarlieva S, Donald CL, Gestuveo RJ, Anaya J-M, Merits A, Burchmore R, Kohl A, Varjak M. 2020. Glucose-Regulated Protein 78 Interacts with Zika Virus Envelope Protein and Contributes to a Productive Infection. Viruses 12:524.

54. Turpin J, Frumence E, Harrabi W, Haddad JG, El Kalamouni C, Desprès P, Krejbich-Trotot P, Viranaïcken W. 2020. Zika virus subversion of chaperone GRP78/BiP expression in A549 cells during UPR activation. Biochimie 175:99–105.

55. Khongwichit S, Sornjai W, Jitobaom K, Greenwood M, Greenwood MP, Hitakarun A, Wikan N, Murphy D, Smith DR. 2021. A functional interaction between GRP78 and Zika virus E protein. Scientific Reports 11:393.

56. Denolly S, Guo H, Martens M, Płaszczyca A, Scaturro P, Prasad V, Kongmanas K, Punyadee N, Songjaeng A, Mairiang D, Pichlmair A, Avirutnan P, Bartenschlager R. 2023. Dengue virus NS1 secretion is regulated via importin-subunit β1 controlling expression of the chaperone GRp78 and targeted by the clinical drug ivermectin. mBio 14:e01441–23.

57. Sornjai W, Promma P, Priewkhiew S, Ramphan S, Jaratsittisin J, Jinagool P, Wikan N, Greenwood M, Murphy D, Smith DR. 2024. The interaction of GRP78 and Zika virus E and NS1 proteins occurs in a chaperone-client manner. Scientific Reports 14:10407.

58. Shrimal S, Cherepanova NA, Gilmore R. 2015. Cotranslational and posttranslocational N-glycosylation of proteins in the endoplasmic reticulum. Semin Cell Dev Biol 41:71–8.

59. Liu T, Hanners NW, Tao H, Szabo C, Xu D, Lin W, Schoggins JW, Yan N, Wu J. 2025. STT3A-mediated mega protein complex assembly during dengue and Zika virus infection. iScience 28:112535.

60. Watterson D, Modhiran N, Young PR. 2016. The many faces of the flavivirus NS1 protein offer a multitude of options for inhibitor design. Antiviral Research 130:7–18.

61. Blight KJ, McKeating JA, Rice CM. 2002. Highly Permissive Cell Lines for Subgenomic and Genomic Hepatitis C Virus RNA Replication. Journal of Virology 76:13001–13014.

62. Eyre NS, Hampton-Smith RJ, Aloia AL, Eddes JS, Simpson KJ, Hoffmann P, Beard MR. 2016. Phosphorylation of NS5A Serine-235 is essential to hepatitis C virus RNA replication and normal replication compartment formation. Virology 491:27–44.

63. Clark DC, Lobigs M, Lee E, Howard MJ, Clark K, Blitvich BJ, Hall RA. 2007. In situ reactions of monoclonal antibodies with a viable mutant of Murray Valley encephalitis virus reveal an absence of dimeric NS1 protein. J Gen Virol 88:1175–1183.

64. Eyre NS, Kirby EN, Anfiteatro DR, Bracho G, Russo AG, White PA, Aloia AL, Beard MR. 2020. Identification of Estrogen Receptor Modulators as Inhibitors of Flavivirus Infection. Antimicrobial Agents and Chemotherapy 64:10.1128/aac.00289-20.

65. Fischl W, Bartenschlager R. 2013. High-Throughput Screening Using Dengue Virus Reporter Genomes, p 205–219. In Gong EY (ed), Antiviral Methods and Protocols doi:10.1007/978-1-62703-484-5_17. Humana Press, Totowa, NJ.

66. Wickham H, Averick M, Bryan J, Chang W, McGowan L, François R, Grolemund G, Hayes A, Henry L, Hester J, Kuhn M, Pedersen T, Miller E, Bache S, Müller K, Ooms J, Robinson D, Seidel D, Spinu V, Yutani H. 2019. Welcome to the Tidyverse. Journal of Open Source Software 4:1686.

67. R. Core Team. 2024. R: A Language and Environment for Statistical Computing.

68. Voß H, Schlumbohm S, Barwikowski P, Wurlitzer M, Dottermusch M, Neumann P, Schlüter H, Neumann JE, Krisp C. 2022. HarmonizR enables data harmonization across independent proteomic datasets with appropriate handling of missing values. Nature Communications 13:3523.

69. Ritchie ME, Phipson B, Wu D, Hu Y, Law CW, Shi W, Smyth GK. 2015. limma powers differential expression analyses for RNA-sequencing and microarray studies. Nucleic Acids Res 43:e47.

70. Kassambara A. 2019. rstatix: Pipe-Friendly Framework for Basic Statistical Tests., vR package version 0.7.2. https://cran.r-project.org/web/packages/rstatix.

71. Wickham H. 2016. ggplot2: Elegant Graphics for Data Analysis. Springer-Verlag New York.

72. Korotkevich G, Sukhov V, Budin N, Shpak B, Artyomov MN. 2021. Fast gene set enrichment analysis. bioRxiv 1.26.0.

73. Eyre NS, Drummer HE, Beard MR. 2010. The SR-BI Partner PDZK1 Facilitates Hepatitis C Virus Entry. PLOS Pathogens 6:e1001130.

